# Beyond Terrestrial Carnivores: Olfactory Bulb Anatomy in a Semi-Aquatic Mammal

**DOI:** 10.64898/2026.11.04.764355

**Authors:** Silvia Alejandra Fraga-Abelleira, Mateo V. Torres, Rocío Canales, Sara Ruiz-Rubio, Pablo Sanchez-Quinteiro, Irene Ortiz-Leal

## Abstract

The organization of the olfactory systems remains poorly understood in mustelids, particularly in lutrines adapted to semiaquatic environments. This study provides the first detailed anatomical, histological, lectin-histochemical, and immunohistochemical characterization of the main (MOB) and accessory (AOB) olfactory bulbs of the Asian small-clawed otter (*Aonyx cinereus*). Serial transverse and sagittal sections were examined using Nissl and Tolivia staining, a panel of lectins, and neuronal and olfactory immunohistochemical markers. Macroscopically, the olfactory bulbs were prominent and elongated, although no distinct AOB could be identified. Histologically, the MOB displayed a well-developed laminar organization, abundant mitral and tufted cells, and a clearly defined lateral olfactory tract. By contrast, the AOB was small, located dorsocaudally within the olfactory peduncle, and lacked the canonical lamination observed in many other mammals. Its identification was supported by the selective binding of Ulex europaeus agglutinin and strong Gαi2 immunoreactivity in glomerular-like structures, whereas Gαo labeling was weak. LEA and STA labeled the superficial layers of both bulbs, while VVA and SBA recognized only scattered MOB glomeruli. Neuronal markers, including NCAM, NSE, NeuN, PGP 9.5, and β-tubulin III, confirmed the neuronal nature of the AOB, while calretinin strongly labeled its glomerular structures and projection neurons. These findings reveal a marked contrast between a highly developed MOB and a small, poorly laminated but neurochemically differentiated AOB. This organization indicates that the vomeronasal pathway is retained in A. cinereus, although in a structurally reduced form, and provides a comparative basis for investigating chemosensory evolution in semiaquatic carnivorans.

## Introduction

The main and accessory olfactory systems play a fundamental role in how mammals interact with their environment. Together, they detect volatile and non-volatile chemical cues involved in numerous biological processes, including foraging, intra- and interspecific recognition, reproduction, and predator detection (Chamero et al. 2007; Bakker and Leinders-Zufall 2016; Torres et al. 2023b). From an anatomical perspective, the processing of chemosensory information has traditionally been attributed to two distinct systems: the main olfactory system (MOS), comprising the olfactory epithelium and main olfactory bulb (MOB) (Firestein 2001; Baum and Cherry 2015), and the accessory olfactory, or vomeronasal, system (VNS), comprising the vomeronasal organ (VNO) and accessory olfactory bulb (AOB) (Halpern and Martínez-Marcos 2003; Larriva-Sahd 2008). The main olfactory system detects a broad range of odorant molecules, particularly, although not exclusively, volatile compounds (Ache and Young 2005; Overath et al. 2014). By contrast, the vomeronasal system has traditionally been associated with the detection of chemical cues involved in social and reproductive behaviour and in interspecific interactions, including pheromonal and kairomonal signals (Dulac and Torello 2003; Fortes-Marco et al. 2013; Tirindelli 2021).

The anatomical organization of these systems reflects their functional specialization. Olfactory sensory neurons located in the olfactory epithelium project their axons to the MOB, where they form synaptic contacts with mitral and tufted cells within the glomeruli (Ortiz-Leal et al. 2022a; Tufo et al. 2022). In parallel, vomeronasal sensory neurons project through the vomeronasal nerves to the AOB (Nakajima et al. 2013; Torres et al. 2022), which represents the first central relay for information processed by the vomeronasal system (Meisami and Bhatnagar 1998; Holy 2018). Although anatomically contiguous with the MOB, the AOB receives a distinct and topographically segregated pattern of primary sensory projections (Prince et al. 2009; Katreddi and Forni 2021). In rodents, lagomorpha and marsupials, the vomeronasal pathway also exhibits a marked neurochemical subdivision (Salazar and Sanchez-Quinteiro 1998; Schneider et al. 2012; Ruiz-Rubio et al. 2024). Apical and basal populations of vomeronasal sensory neurons are associated with Gαi2- and Gαo-mediated signalling pathways, respectively, and project to the anterior and posterior regions of the AOB (Halpern et al. 1995; Takigami 2004; Kondoh et al. 2025b). Nevertheless, the traditional view of the main and accessory olfactory systems as entirely separate pathways has increasingly been questioned. Anatomical and functional interactions between them have been demonstrated, indicating that both systems may act in a coordinated manner during certain chemically guided behaviours (Baum and Cherry 2015; Salazar et al. 2016).

The vomeronasal system is particularly relevant to intraspecific chemical communication. It detects cues capable of modulating reproductive behaviour, social recognition, aggression, territoriality, and the establishment of social hierarchies (Estes 1972; Kelliher 2007; Pallé et al. 2020; Asproni et al. 2022). It is also involved in neuroendocrine responses associated with reproductive and maternal behaviours (Beltramino and Taleisnik 2008; Numan 2026; Sanahuja-Irene et al. 2026). Although much of the current knowledge in this field derives from laboratory rodents, the organization and degree of development of the vomeronasal system vary widely among mammals at both morphological and molecular levels (Salazar et al. 1998a; Salazar et al. 2007; Nikaido 2019) . Such diversity limits the extent to which mechanisms characterized in experimental rodent models, particularly mice, can be extrapolated to other mammalian species. Comparative studies are therefore essential for understanding the evolution and functional specialization of the vomeronasal system across different mammalian lineages (Salazar and Sanchez Quinteiro 2009; Kondoh et al. 2020).

Among mammals, carnivorans display substantial diversity in the organization of the VNS. Although the VNO and AOB have been identified in numerous carnivoran species, their degree of development, anatomical organization, and neurochemical characteristics differ considerably among families and even between closely related species (Salazar and Sánchez-Quinteiro 2011; Tomiyasu et al. 2017; Torres et al. 2021; Ortiz-Leal et al. 2024; Ortiz-Leal et al. 2024). Canids provide a particularly illustrative example. In the domestic dog, the AOB exhibits a relatively poorly differentiated organization (Nakajima et al. 1998; Salazar et al. 2013), whereas recent studies of wild canids have revealed a greater structural complexity. The red fox, for example, possesses a well-developed VNO and a clearly differentiated AOB whose organization contrasts with that reported in the domestic dog (Ortiz-Leal et al. 2020; Ortiz-Leal et al. 2023). These differences have prompted the hypothesis that evolutionary changes associated with domestication may have contributed to the reduction or reorganization of certain components of the vomeronasal system in domestic canids (Ortiz-Leal et al. 2022b). More broadly, the degree of development of this system may reflect not only phylogenetic constraints but also ecological, evolutionary, and behavioural factors.

Mustelids constitute another carnivoran group of particular interest because relatively little is known about the organization of their vomeronasal system. Studies in mink (*Mustela vison*) demonstrated the presence of a well-defined VNO, including sensory epithelium and structural and vascular specializations characteristic of the VNO (Salazar et al. 1994a; Salazar et al. 1995). Subsequent lectin-histochemical studies of the mink olfactory bulb allowed the presumptive AOB to be identified, although its organization differed from that described in other mammalian species (Salazar et al. 1998b). In the domestic ferret (*Mustela putorius furo*), both the VNO and AOB are present, although the AOB is relatively small, displays an indistinct laminar organization, and is difficult to delineate using conventional histological techniques and marker (Weiler 1999; Kelliher et al. 2001; Woodley et al. 2004). These characteristics may explain why previous studies described the AOB of some mustelids as absent, rudimentary, or poorly defined (Jawlowski 1956). This observations highlight the importance of combining conventional histology with complementary immunohistochemical and lectin-histochemical approaches to identify this structure reliably (Salazar et al. 1998b; Kelliher et al. 2001).

Within Mustelidae, further investigation of this variability is particularly relevant in the subfamily Lutrinae, which comprises species adapted to aquatic and semiaquatic environments. The Asian small-clawed otter (*Aonyx cinereus*) is a small-bodied lutrine inhabiting rivers, wetlands, and rice fields (Hussain et al. 2011). Its complex social behaviour raises specific questions concerning the role of chemosensory communication. In aquatic environments, the availability, persistence, and dispersal of chemical cues differ from those prevailing in terrestrial habitats (Catania 2006; Mollo et al. 2014; Mollo et al. 2017). This species therefore provides an interesting model for investigating whether and how the main and accessory olfactory systems have been modified in association with a semiaquatic lifestyle.

In cetaceans, adaptation to a fully aquatic environment has profoundly affected the main olfactory system, although its degree of regression differs between the two major lineages. Odontocetes, including dolphins and toothed whales, have lost the olfactory bulbs and associated olfactory pathways, whereas mysticetes retain a reduced but anatomically recognizable main olfactory system and a limited repertoire of functional olfactory receptor genes (Kishida et al. 2015b; Kishida et al. 2015a; De Vreese et al. 2023). By contrast, the VNO and its associated accessory olfactory pathways are considered absent in cetaceans (Thewissen et al. 2011; Berta et al. 2014; De Vreese et al. 2023).

Among pinnipeds -semiaquatic marine carnivorans-the available evidence reveals a more variable pattern. In the harbour seal (*Phoca vitulina*; Phocidae), neither a sensory VNO nor a recognizable AOB has been identified, suggesting the loss of the vomeronasal pathway (Kondoh et al. 2024). By contrast, a recent study demonstrated the presence of an anatomically and neurochemically differentiated vomeronasal system in the Steller sea lion (*Eumetopias jubatus*; Otariidae), indicating the retention of a functional V1R–Gαi2 pathway (Kondoh et al. 2025a).

The preservation of the vomeronasal system in other semiaquatic mammals further demonstrates that adaptation to aquatic environments does not necessarily entail the loss of this sensory pathway. A differentiated VNO and AOB have been described in the capybara (Torres et al. 2020). Similarly, both structures are well developed in the Eurasian beaver, in which the vomeronasal system has been proposed to participate in the detection of non-volatile compounds involved in social communication, particularly those derived from glandular secretions deposited by conspecifics (Tomiyasu et al. 2022). Taken together, these findings indicate that the retention, reduction, or loss of the vomeronasal system among aquatic and semiaquatic mammals does not follow a uniform pattern and may reflect lineage-specific ecological and behavioural pressures.

Genomic studies further suggest that the transition to aquatic and semiaquatic environments may have influenced chemosensory function in some lutrines. Extensive pseudogenization of olfactory receptor genes has been reported in the sea otter (*Enhydra lutris*) and giant otter (*Pteronura brasiliensis*) (Beichman et al. 2019), while *TRPC2*, a key component of vomeronasal signal transduction, has been described as a pseudogene in the North American river otter (*Lontra canadensis*) (Yu et al. 2010; Beichman et al. 2019). However, these findings are restricted to a small number of species and cannot be generalized to Lutrinae as a whole. However, the anatomical and neurochemical correlates of these genomic changes remain largely unknown, as the detailed organization of the vomeronasal system in lutrines has yet to be characterized. To our knowledge, no previous study has specifically characterized the morphology, histology, or neurochemical features of the AOB in lutrines. Information concerning the organization of their MOB is likewise scarce. A detailed characterization of these structures is therefore required to determine whether the olfactory and vomeronasal systems of *A. cinereus* conform to the general pattern described in other mustelids or exhibit distinctive features potentially associated with its semiaquatic ecology.

In this context, the present study provides an anatomical, histological, immunohistochemical, and lectin-histochemical characterization of the structures comprising the main and accessory olfactory systems of A. cinereus, with particular emphasis on the identification and organization of the MOB and AOB. The immunohistochemical analysis included the neuronal markers calbindin and calretinin, whereas the lectin-histochemical study employed UEA, LEA, and SBA. The combined use of these approaches enabled an integrated assessment of the structural and neurochemical organization of the olfactory bulbs.

## Methods

### Animals and tissue collection

The study was performed on the olfactory bulbs of two adult female Asian small-clawed otters (*Aonyx cinereus*) housed at Mundomar Benidorm (Alicante, Spain). Both animals died from causes unrelated to the present study, and no animal was euthanized or subjected to any experimental procedure for research purposes. Immediately after death, the skin, eyes, cranial musculature, mandible, and dorsal portion of the skull were removed to facilitate penetration of the fixative. The heads were immersed in 10% formalin and transferred to the Faculty of Veterinary Medicine, University of Santiago de Compostela, Lugo, Spain, under the framework of a collaboration agreement between the zoological institution and the Animal Morphology Research Group. Because the study was performed exclusively on post-mortem material obtained from animals that had died independently of the investigation, specific approval for animal experimentation was not required under Directive 2010/63/EU. The collection, transfer, and scientific use of the specimens were authorized by the zoological institution.

### Brain extraction and macroscopic examination

The remaining lateral walls and base of the neurocranium were carefully removed using rongeurs. The dura mater was opened and reflected laterally, and the brain was extracted with particular care to preserve the olfactory bulbs and peduncles. After extraction, the brain was examined and photographed in dorsal, lateral, medial, and ventral views. The size, shape, orientation, and topographical relationships of the olfactory bulbs, olfactory peduncles, lateral olfactory tracts, olfactory tubercles, and piriform lobes were recorded. The olfactory bulbs were subsequently separated from the remainder of the brain at the level of the olfactory peduncle and processed for histological, lectin-histochemical, and immunohistochemical examination.

### Tissue processing and sectioning

Following fixation, the olfactory bulbs were washed in distilled water, dehydrated through a graded ethanol series, cleared in xylene, and infiltrated with paraffin wax using an automated tissue processor. The specimens were then embedded individually in paraffin blocks, with their orientation selected to obtain complete sections of the olfactory bulb and peduncle. Serial sections, 8 µm thick, were obtained using a rotary microtome. For each specimen, one olfactory bulb was sectioned in the transverse plane and the contralateral bulb in the sagittal plane. Sections were mounted on adhesive-coated glass slides and distributed into parallel series for conventional histology, lectin histochemistry, and immunohistochemistry.

### Conventional histology

Selected sections were stained using the Nissl method to examine neuronal cytoarchitecture and identify the cellular layers of the main and accessory olfactory bulbs. Additional sections were processed using the Tolivia method for the simultaneous visualization of neuronal somata and myelinated fiber systems in paraffin-embedded nervous tissue (Tolivia et al. 1988; Salazar et al. 2007).

The stained series were used to assess the general laminar organization of the main olfactory bulb (MOB), delimit the olfactory nerve, glomerular, external plexiform, mitral-cell, internal plexiform, and granule-cell layers, and identify the lateral olfactory tract. Particular attention was paid to the dorsocaudal region of the olfactory bulb and adjacent olfactory peduncle in order to locate the accessory olfactory bulb (AOB) and evaluate its cytoarchitecture and relationship with the surrounding MOB.

### Lectin histochemistry

Lectin histochemical study was performed to characterize the distribution of specific glycoconjugates within the MOB and AOB. Five biotinylated lectins were employed: *Ulex europaeus* agglutinin I (UEA-I), *Lycopersicon esculentum* agglutinin (LEA), *Solanum tuberosum* agglutinin (STA), *Vicia villosa* agglutinin (VVA), and *Glycine max* agglutinin (SBA). Lectin specificity and working conditions are summarized in Table 1. Lectins recognize specific terminal carbohydrate residues and have been extensively used to demonstrate neurochemical heterogeneity within mammalian olfactory pathways (Lis and Sharon 1998).

**Table 1.** Lectins used, concentrations and binding specificities.

| Lectin | Abbrev. | Dilution mg/ml | Supplier / catalogue no. | Principal carbohydrate specificity |
|---|---|---|---|---|
| *Ulex europaeus* agglutinin I | UEA-I | 2.0 | Vector Laboratories<br>B-1065 | α-L-fucose |
| *Lycopersicon esculentum* agglutinin | LEA | 1.0 | Vector Laboratories<br>B-1175 | Poly-N-acetyllactosamine and β1-4-linked N-acetylglucosamine oligomers |
| *Solanum tuberosum* agglutinin | STA | 2.0 | Vector Laboratories<br>B-1165 | N-acetylglucosamine oligomers and N-acetyllactosamine |
| *Vicia villosa* agglutinin | VVA | 2.0 | Vector Laboratories<br>B-1235 | Terminal N-acetylgalactosamine |
| *Glycine max* agglutinin | SBA | 2.0 | Vector Laboratories<br>B-1015 | Terminal α- or β-linked N-acetylgalactosamine and galactose |

UEA-I recognizes terminal α-L-fucosyl residues and has been shown to delineate specific olfactory and vomeronasal projections, including the AOB in rodents and wild canids (Lee et al. 2012; Kondoh et al. 2017; Ortiz-Leal et al. 2022b). LEA and STA bind N-acetylglucosamine-rich glycoconjugates and have been used to visualize sensory axons and glomerular structures in both the MOB and AOB of several mammalian species (Ichikawa et al. 1992). VVA and SBA predominantly recognize terminal N-acetylgalactosamine-containing glycoconjugates; VVA, in particular, has proved useful for identifying vomeronasal axons and regional subdivisions within the AOB of rats, opossums and horses (Shapiro et al. 1995; Chun et al. 2024). Because lectin-binding patterns may vary considerably among species, the complete panel was applied to determine whether the distribution of these glycoconjugates could differentiate accessory from main olfactory territories in *Aonyx cinereus*.

Paraffin sections were deparaffinized in xylene and rehydrated through a descending ethanol series. Endogenous peroxidase activity was blocked by incubation in 3% hydrogen peroxide in distilled water for 10 min. After washing in 0.1 M phosphate buffer (PB; pH 7.2), nonspecific binding sites were blocked with 2% bovine serum albumin for 30 min at room temperature. Sections were incubated overnight at 4 °C with the corresponding biotinylated lectin diluted in PB. After three washes in PB, the sections were incubated for 90 min at room temperature with an avidin–biotin– peroxidase complex (Vectastain ABC Kit, Vector Laboratories, Burlingame, CA, USA). Peroxidase activity was visualized using 0.05% 3,3′-diaminobenzidine tetrahydrochloride (DAB) containing 0.003% hydrogen peroxide in 0.2 M Tris–HCl buffer (pH 7.6). Sections were subsequently washed, dehydrated, cleared in xylene, and coverslipped with a permanent mounting medium. Selected sections were counterstained with hematoxylin.

Negative controls were processed in parallel by omitting the corresponding lectin from the incubation solution.

### Immunohistochemistry

The immunohistochemical study was performed on serial sections to characterize neuronal populations, sensory projections, neuronal maturation, calcium-binding proteins, and the molecular phenotype of the main and accessory olfactory pathways. The primary antibodies employed included neuronal and cytoskeletal markers (NCAM, NSE, NeuN, PGP 9.5, and β-tubulin III), olfactory sensory-neuron markers (OMP), G-protein α subunits associated with vomeronasal signal transduction (Gαi2 and Gαo), and the calcium-binding proteins calretinin, calbindin D-28k, and parvalbumin. Antibody sources, catalogue numbers, host species, and working dilutions are provided in Table 2.

**Table 2.** Detailed information on the primary antibodies used in this study, including host species, working dilution, supplier and catalogue number, and target immunogen.

| Primary antibody | Host species | Dilution | Supplier / Catalogue number | Target immunogen |
|---|---|---|---|---|
| Anti-Gαo | Rabbit | 1:200 | Santa Cruz Biotechnology sc-387 | Bovine GTP-binding protein Gαo subunit |
| Anti-Gαi2 | Rabbit | 1:100 | Santa Cruz Biotechnology sc-7276 | Peptide mapping within a highly divergent domain of rat Gαi2 |
| Anti-OMP | Mouse | 1:100 | Santa Cruz Biotechnology sc-365818 | Amino acids 1–163 of human olfactory marker protein |
| Anti-NCAM | Mouse | 1:100 | Santa Cruz Biotechnology sc-59864 | Human neural cell adhesion molecule |
| Anti-NSE | Rabbit | 1:150 | Cloud-Clone Corp., PAA537Mu01 | Recombinant mouse neuron-specific enolase expressed in *Escherichia coli* |
| Anti-NeuN | Mouse | 1:100 | Merck Millipore, MAB377, clone A60 | Purified neuronal nuclei from mouse brain |
| Anti-PGP 9.5 | Rabbit | 1:200 | Proteintech 14730-1-AP | UCHL1/PGP 9.5 fusion protein Ag6490 |
| Anti-βIII-tubulin | Rabbit | 1:300 | Proteintech 10068-1-AP | βIII-tubulin fusion protein Ag0117 |
| Anti-calretinin | Rabbit | 1:200 | Proteintech 12278-1-AP | Recombinant human calretinin containing an N-terminal 6×His tag |
| Anti-calbindin D-28k | Rabbit | 1:200 | Proteintech 14479-1-AP | Recombinant rat calbindin D-28k expressed in *Escherichia coli* |
| Anti-parvalbumin | Rabbit | 1:100 | Proteintech 26521-1-AP | Recombinant parvalbumin-derived peptide |

Abbreviations: CB, calbindin D-28k; CR, calretinin; Gαi2, G-protein αi2 subunit; Gαo, G-protein αo subunit; NCAM, neural cell adhesion molecule; NSE, neuron-specific enolase; NeuN, neuronal nuclear protein; OMP, olfactory marker protein; PGP 9.5, protein gene product 9.5/ubiquitin C-terminal hydrolase L1; PV, parvalbumin; βIII-tubulin, class III β-tubulin.

Before overnight incubation with the primary antibodies, sections were processed following the same procedure described above for lectin histochemistry, including deparaffinization, rehydration, inhibition of endogenous peroxidase activity), and blocking of nonspecific binding with 2% BSA. Sections were subsequently incubated overnight at 4 °C with the corresponding primary antibodies at the dilutions specified in Table 2. After washing in PB, antibody binding was detected using the appropriate species-specific ImmPRESS HRP polymer reagent (anti-rabbit or anti-mouse IgG; Vector Laboratories) for 30 min at room temperature. Immunoreactivity was visualized with 0.05% DAB and 0.003% hydrogen peroxide in 0.2 M Tris–HCl buffer (pH 7.6). Sections were then washed, dehydrated, cleared, and coverslipped. Selected sections were lightly counterstained with hematoxylin to facilitate identification of cellular and laminar boundaries.

For each marker, negative-control sections were processed under identical conditions, except that the primary antibody was omitted. Sections from tissues previously known to express the corresponding antigen were used as positive controls when available.

### Microscopic examination and image acquisition

Sections were examined using a Olympus B50 light microscope equipped with a digital camera. Images were acquired using cellSens software under standardized illumination and exposure conditions. Only global adjustments of brightness, contrast, and color balance were applied, and these were performed uniformly across the entire image. The identification of the MOB layers was based on their cytoarchitecture, cellular composition, and topographical relationships. Because the AOB was not macroscopically distinguishable, its identification was based on the combined assessment of its dorsocaudal and sagittal location, cytoarchitectural organization, lectin-binding profile, and immunoreactivity for vomeronasal and neuronal markers. Given the descriptive nature of the study and the limited availability of post-mortem specimens of this species, no inferential statistical analyses were performed. Findings were considered consistent when the same anatomical or neurochemical pattern was repeatedly observed in corresponding regions across serial sections and corroborated by complementary histological, lectin-histochemical, or immunohistochemical markers.

## Results

### Macroscopical study

Macroscopic examination of the brain of *Aonyx cinereus* revealed a marked development of the olfactory system. In lateral view, the olfactory bulb (OB) appears as a prominent structure with an elongated and pointed morphology, clearly projecting rostrally relative to the telencephalon, giving the anterior pole of the brain a characteristic elongated appearance (Fig. 1A). In the ventral view of the brain, the principal elements of the main olfactory system were clearly identifiable, including the OB, olfactory peduncles, olfactory tubercle, and piriform lobe, all showing a notable degree of development (Fig. 1B). The olfactory peduncles exhibit an elongated configuration and contain the olfactory tract, providing the connection between the OB and more caudal telencephalic regions. Detailed inspection of the OB in both lateral and medial views did not reveal, at the macroscopic level, any clearly distinguishable structure that could be unequivocally attributed to an AOB (Fig. 1C). However, based on subsequent analysis of the histological series, a specific area within the OB was delineated as the presumptive location of this component. This region, indicated in the figure 1C, does not present distinctive morphological features under macroscopic observation.

**Figure 1.**
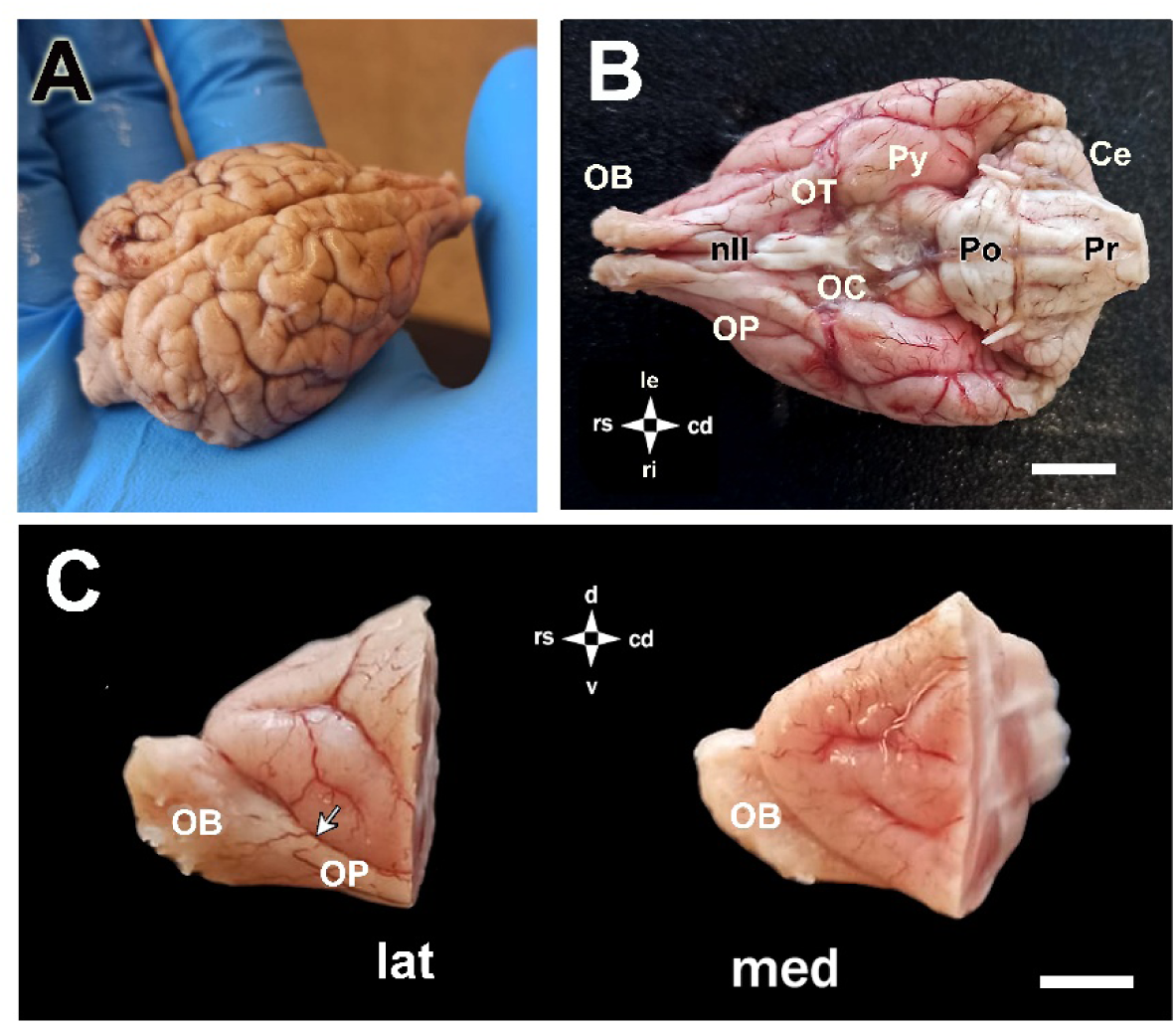
**Macroscopic study of the olfactory system of the Asian small-clawed otter (*Aonyx cinereus*)**. **(A)** Lateral view of the brain from one of the processed specimens, showing a elongated, and pointed olfactory bulb (OB), clearly projecting rostrally, together with the rostral portion of the telencephalon. (**B)** Ventral view of the brain, in which the main components of the olfactory system can be identified, including the OB, olfactory peduncle (OP), olfactory tubercle (OP), and piriform lobe (Py), all of which are notably well developed in this species. The olfactory peduncles contain the white matter of the olfactory tract. **(C)** Lateral (lat) and medial (med) views of the forebrain. The arrow indicates the area that, based on subsequent microscopic histological analysis, is considered representative of the presumptive location of the AOB. At the macroscopic level, no distinct structural features corresponding to this component can be identified in this region. Abbreviations: Ce: Cerebellum; nII: Optic nerves; OC: Optic chiasm; Po: Pons; Pr: Pyramids. Scale bar: B = 1 cm; C = 500 μm.

### Histological organization of the main and accessory olfactory bulbs

For the microscopic analysis of the olfactory bulbs of *Aonyx cinereus*, Nissl staining was applied to serial sections obtained in both transverse and sagittal anatomical planes (Figs. 2-4). This approach allowed a detailed evaluation of the cytoarchitectural organization of the MOB, as well as a systematic search for morphological evidence of the AOB. Serial sections were examined throughout the rostrocaudal extent of the OBs to characterize their laminar organization, cellular density, and neuronal composition of the MOB, and to identify potential regions corresponding to the accessory olfactory system.

**Figure 2.**
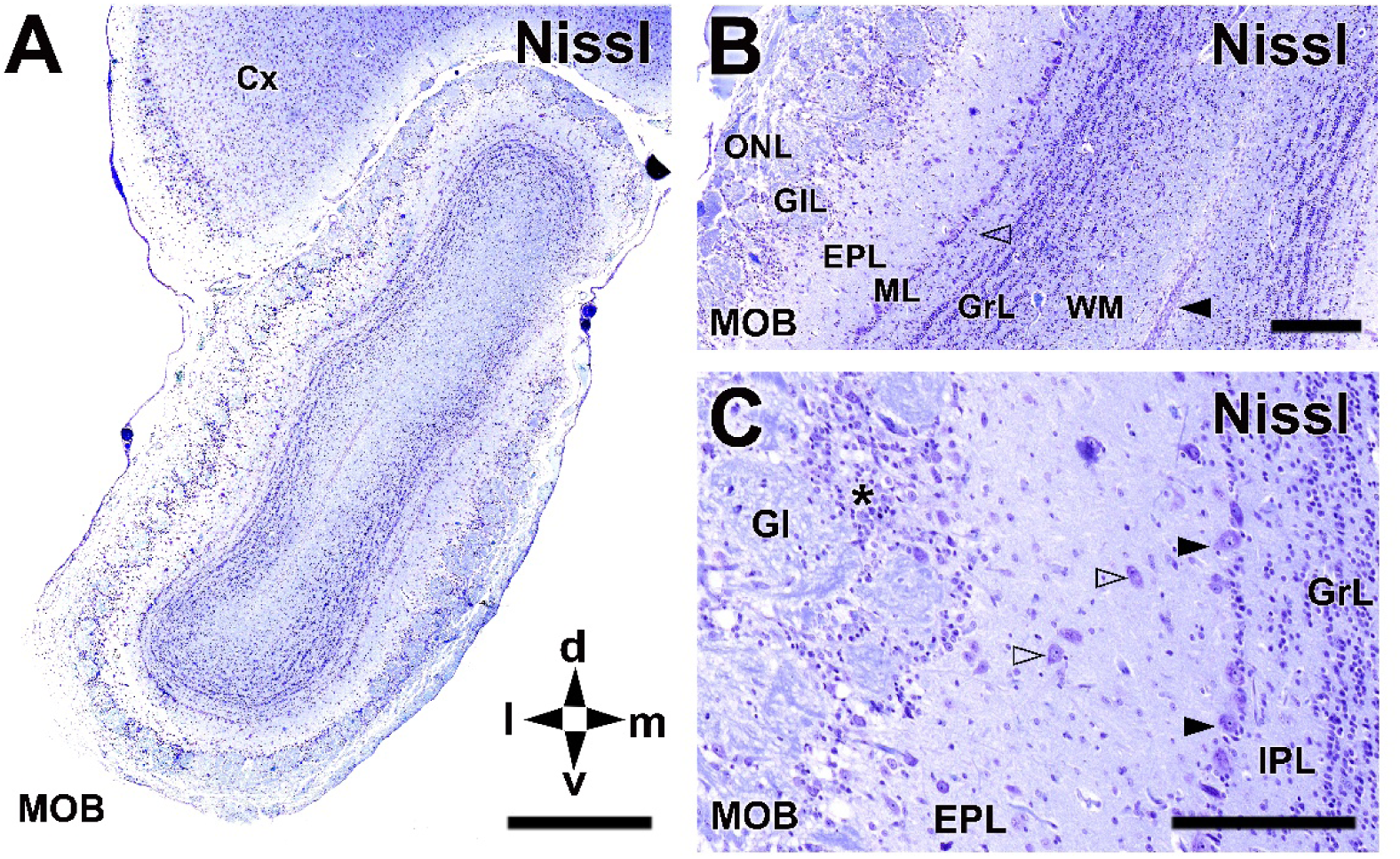
Histological organization of the MOB of the Asian small-clawed otter (*Aonyx cinereus*). Nissl staining. **(A)** Transverse section through a central level of the MOB showing its characteristic laminar organization and overall cytoarchitecture. The orderly stratification of the bulb and the differences in cellular density among layers are clearly distinguishable. **(B)** Higher-magnification view illustrating the distinct layers of the MOB, including the olfactory nerve layer (NL), glomerular layer (GL), external plexiform layer (EPL), mitral cell layer (ML), internal plexiform layer (open arrowhead), granular layer (GrL), and white matter (WM). The arrowhead indicates the subventricular zone. **(C)** High-magnification view of the intermediate layers of the MOB. Huge mitral cells are identifiable within the ML (arrowheads), while numerous tufted cells are observed in the EPL (open arrowheads). A high concentration of periglomerular cells surrounds the glomeruli, particularly in their deeper regions (*). Granule cells are arranged in clusters with a linear organization within the granular layer. Scale bars: A = 1 mm; B,C = 250 µm.

The MOB exhibited a well-defined laminar organization, consistent with that described in other mammalian species. This arrangement was clearly visible in the transverse plane (Fig. 2). At higher magnification, the olfactory nerve layer (NL), glomerular layer (GlL), external plexiform layer (EPL), mitral cell layer (ML), internal plexiform layer (IPL), granular layer (GrL), and underlying white matter (WM) could be readily distinguished (Fig. 2B). The subventricular zone was identifiable adjacent to the white matter, indicating ongoing neurogenic potential within the bulb. Detailed examination of the intermediate layers demonstrated a prominent development of mitral cells within the mitral cell layer (Fig. 2C). Numerous tufted cells were observed within the external plexiform layer, and around the glomeruli, a high concentration of periglomerular cells was evident, particularly in the deeper portions of the glomerular layer. Granule cells in the granular layer were arranged in well-defined clusters, displaying a linear organization.

Serial histological examination of the entire OB complex, from its most rostral extent to its caudoventral termination at the level of the olfactory peduncle, did not reveal any structure corresponding to the canonical organization of the AOB as described in other mammalian groups, including rodents, lagomorphs, ruminants, or felids among carnivores. In none of the sections examined was a well-defined AOB with the typical laminar arrangement—comprising a distinct glomerular layer, mitral-plexiform cell layer, and granule cell layer—observed. Instead, only a limited number of glomerular-like structures were observed, displaced to a dorsomedial position in the caudal portion of the olfactory peduncle (Fig. 3A, B). These structures displayed a loose and poorly organized appearance. At higher magnification, the presumptive AOB region showed an atypical and rudimentary cytoarchitecture, characterized by the absence of clear lamination and by a heterogeneous mixture of neuropil and neuronal elements (Fig. 3C, D). Neurons with large nuclei and prominent central nucleoli, likely corresponding to projection neurons, were interspersed with small-sized cells consistent with granule neurons. This atypical organization was consistently observed in all the olfactory bulbs analyzed.

**Figure 3.**
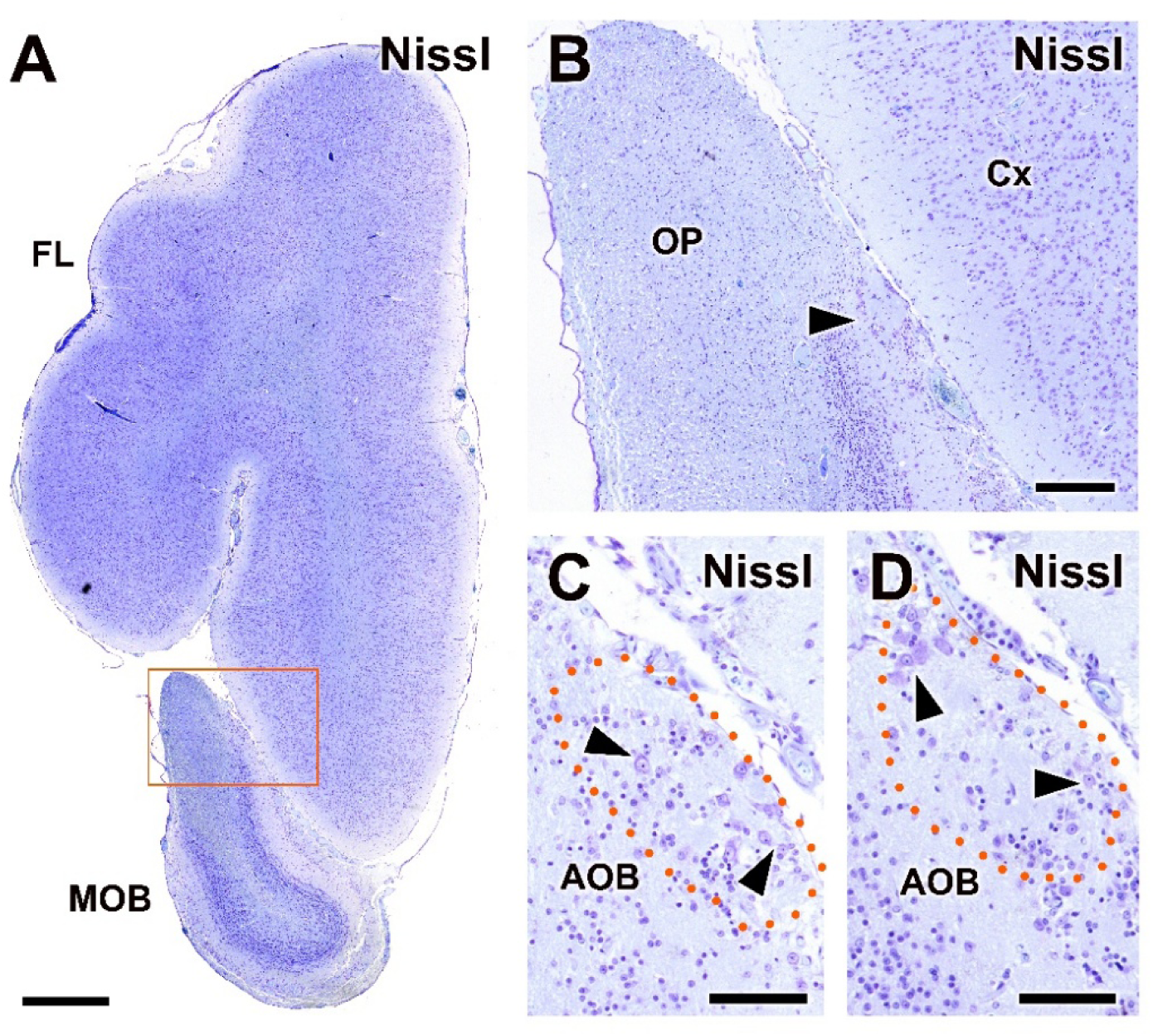
Transverse histological study of the OB of the Asian small-clawed otter (*Aonyx cinereus*). Nissl staining. **(A)** Transverse section through the rostral portion of the brain at the level of the olfactory peduncle and frontal lobe (FL), including the ventro caudal part of the MOB. The topographical distribution of the presumptive AOB is indicated. The boxed area is shown at higher magnification in (B). **(B)** Higher-magnification view showing the presumptive AOB (arrowhead) located dorsomedially to the caudal MOB, within the olfactory peduncle and in close topographical relationship with the cerebral cortex (Cx). **(C, D)** Transverse sections of the presumptive AOB. A rudimentary organization is observed, characterized by alternating regions of neuropil and neuronal somata. Neurons with large nuclei and visible nucleoli (arrowheads), likely corresponding to projection neurons, are intermingled with small-sized cells consistent with granule cells. The dotted line outlines the presumptive AOB region. Scale bars: A = 1 mm; B = 250 µm; C,D = 100 µm.

**Figure 4.**
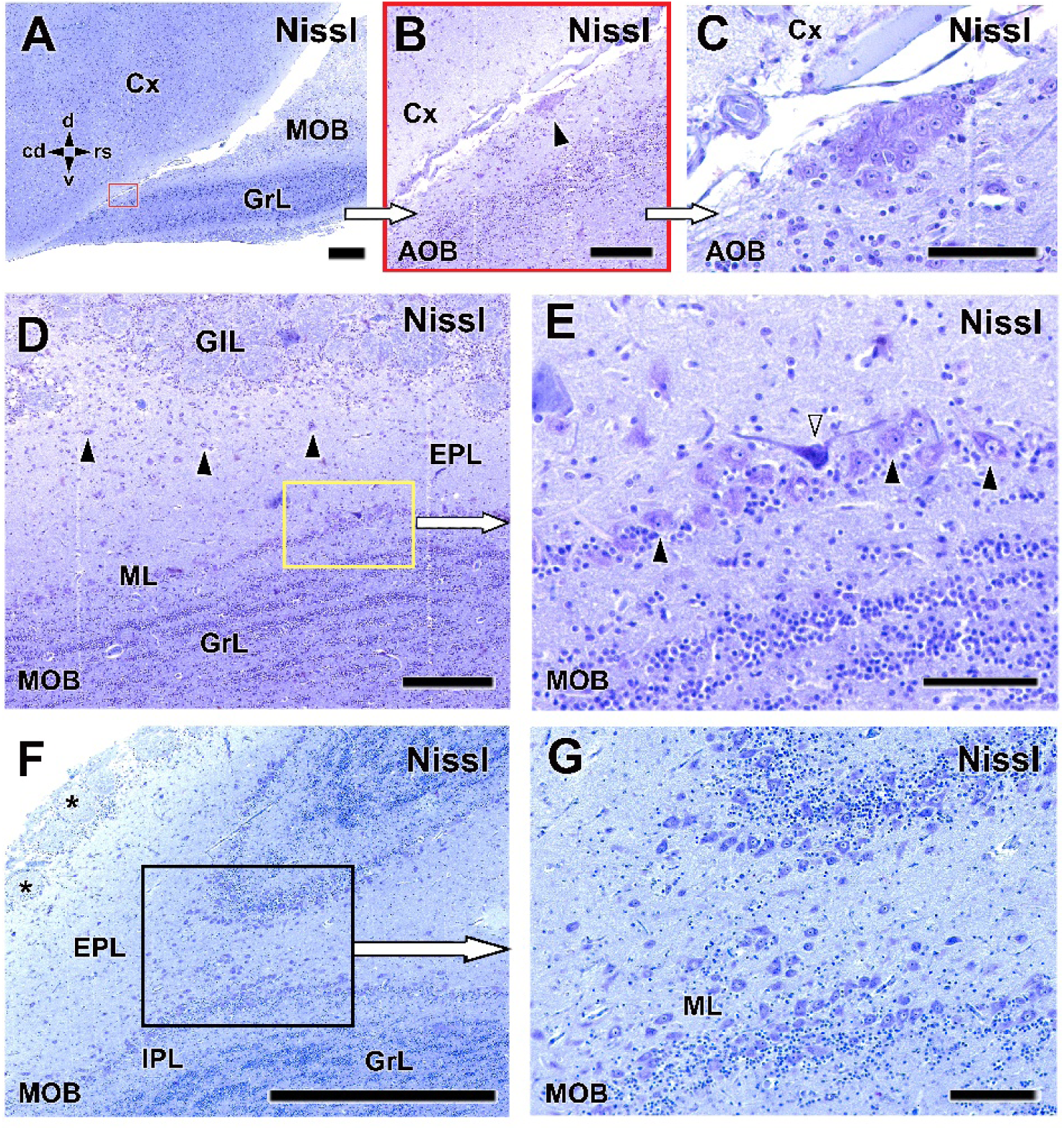
Sagittal histological study of the main and accessory OB of the Asian small-clawed otter (*Aonyx cinereus*). Nissl staining. (A–C) Sagittal sections of the caudal portion of the MOB, allowing the identification of the presumptive AOB. In (A), the presumptive AOB is located in a caudodorsal position within the olfactory peduncle. Higher magnification in (B) reveals an undifferentiated structure which, at higher magnification (C), appears to be partially composed of neurons with large somata and conspicuous nucleoli, centrally positioned. This atypical neuronal aggregation corresponds to the presumptive AOB. **(D,E)** Sagittal sections through the central region of the MOB, showing its characteristic laminar organization. The internal plexiform layer is well defined, and the mitral cell layer is composed of dense alignments of mitral cells (arrowheads). In some cases, mitral cells are accompanied by deep bipolar interneurons with short axons (open arrowheads). **(F,G)** Medial parasagittal sections of the MOB. The tangential orientation of the section plane allows appreciation of the abundance and morphological diversity of mitral and tufted cells characteristic of this semi-aquatic species. Scale bars: A,F = 1 mm; G = 500 µm; B-D = 250 µm; C,E = 100 µm.

Serial sagittal histological sections provided complementary information to the transverse analysis of the OB complex. Examination of the caudal region of the main olfactory bulb at the level of the olfactory peduncle allowed the identification of the presumptive AOB, located in a dorsocaudal position (Fig. 4A–C). At low magnification, this structure appears poorly differentiated; however, higher magnification reveals a compact neuronal aggregation composed predominantly of neurons with large somata and conspicuous nucleoli, centrally arranged. Analysis of the MOB in sagittal sections allowed a detailed characterization of its laminar organization. In central sagittal sections, the typical stratification of the MOB was clearly identifiable, including a well-defined IPL and a ML formed by dense alignments of mitral cells (Fig. 4D, E). In some instances, these mitral cells were accompanied by deep bipolar interneurons with short axons. In medial sagittal sections, the tangential orientation of the plane of section facilitated the observation of both the abundance and the morphological diversity of mitral and tufted cells within the main olfactory bulb (Fig. 4F, G). The variety of soma sizes, shapes, and dendritic profiles observed is consistent with the complex organization of the olfactory circuitry in this semi-aquatic species.

Tolivia staining allowed a clear visualization of myelinated fibers and neuronal cell bodies within the OB. Myelinated fibers appeared intensely stained in black, whereas neurons and interneurons were stained in red. At low magnification, a strong staining corresponding to myelinated structures was observed in the internal capsule of the telencephalon and in the deep layers of the MOB (Fig. 5A). At higher magnification, the laminar organization of the olfactory bulb was well preserved, enabling the identification of the external plexiform layer, where mitral and tufted cells were clearly visible (Fig. 5B). Glomeruli were sharply outlined by periglomerular cells, while myelinated fibers were largely confined to the granular layer. Notably, a dense and well-defined bundle of myelinated fibers forming the lateral olfactory tract (LOT) was observed extending from the MOB, indicating the effectiveness of Tolivia staining for visualizing major olfactory pathways (Fig. 5C).

**Figure 5.**
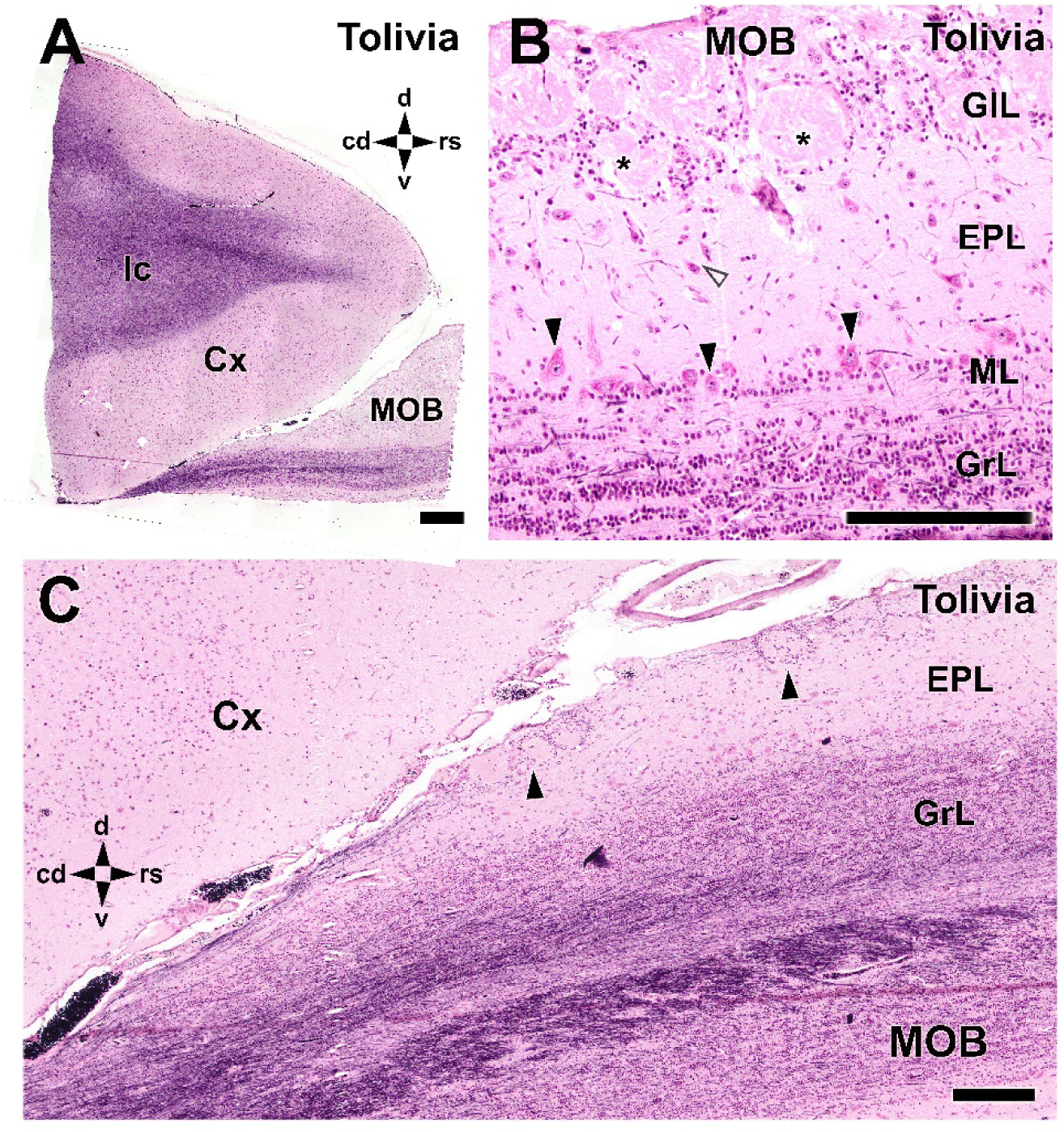
**Tolivia staining of the MOB**. **(A)** Low-magnification view of the rostral portion of the encephalon showing intense staining in the internal capsule of the frontal lobe and in the deep regions of the MOB, indicative of a high density of myelinated fibers. **(B)** Higher magnification of the MOB revealing its characteristic lamination. Mitral (arrowhead) and tufted cells (open arrowhead) are clearly identified within the EPL, and glomeruli (asterisk) are well delineated by surrounding periglomerular cells. Myelinated fibers are mainly restricted to the granular layer (GrL). **(C)** A prominent bundle of myelinated fibers forming the lateral olfactory tract (LOT) is evident adjacent to the granular layer and extending caudally from the MOB. Black arrowheads indicate groups of glomeruli located in the caudal and caudodorsal regions of the MOB. Scale bars: A = 1 mm; B,C = 250 µm.

### Lectin histochemical study

Lectins are carbohydrate-binding proteins that have been widely used as histochemical markers in the study of the main and accessory olfactory systems, as some of them allow a differential identification of both pathways. For this reason, we applied a panel of five lectins -UEA, LEA, STA, VVA and SBA- to evaluate their binding patterns within the olfactory bulb and to assess their usefulness in distinguishing the main olfactory bulb from the accessory olfactory bulb.

UEA lectin histochemistry revealed a highly restricted binding pattern within the olfactory bulb. At low magnification, no UEA labeling was detected in the telencephalon or in the MOB, with the exception of a discrete region located at the presumed position of the AOB (Fig. 6A). Higher magnification of this UEA-positive area showed a strongly labeled glomerular-like structure, accompanied by smaller UEA-positive profiles, presumably corresponding to branches of the vomeronasal nerve (Fig. 6B). When sections were counterstained with hematoxylin, the UEA-positive structure was observed to be surrounded by UEA-negative neurons displaying large, rounded nuclei, together with smaller, also UEA-negative cells distributed throughout the adjacent tissue, consistent with the cellular organization of the AOB (Fig. 6C). A broader view including both the accessory and main olfactory bulbs confirmed that UEA labeling was strictly confined to the AOB, with no detectable binding in the MOB (Fig. 6D), supporting the specificity of UEA as a histochemical marker for the accessory olfactory system.

**Figure 6.**
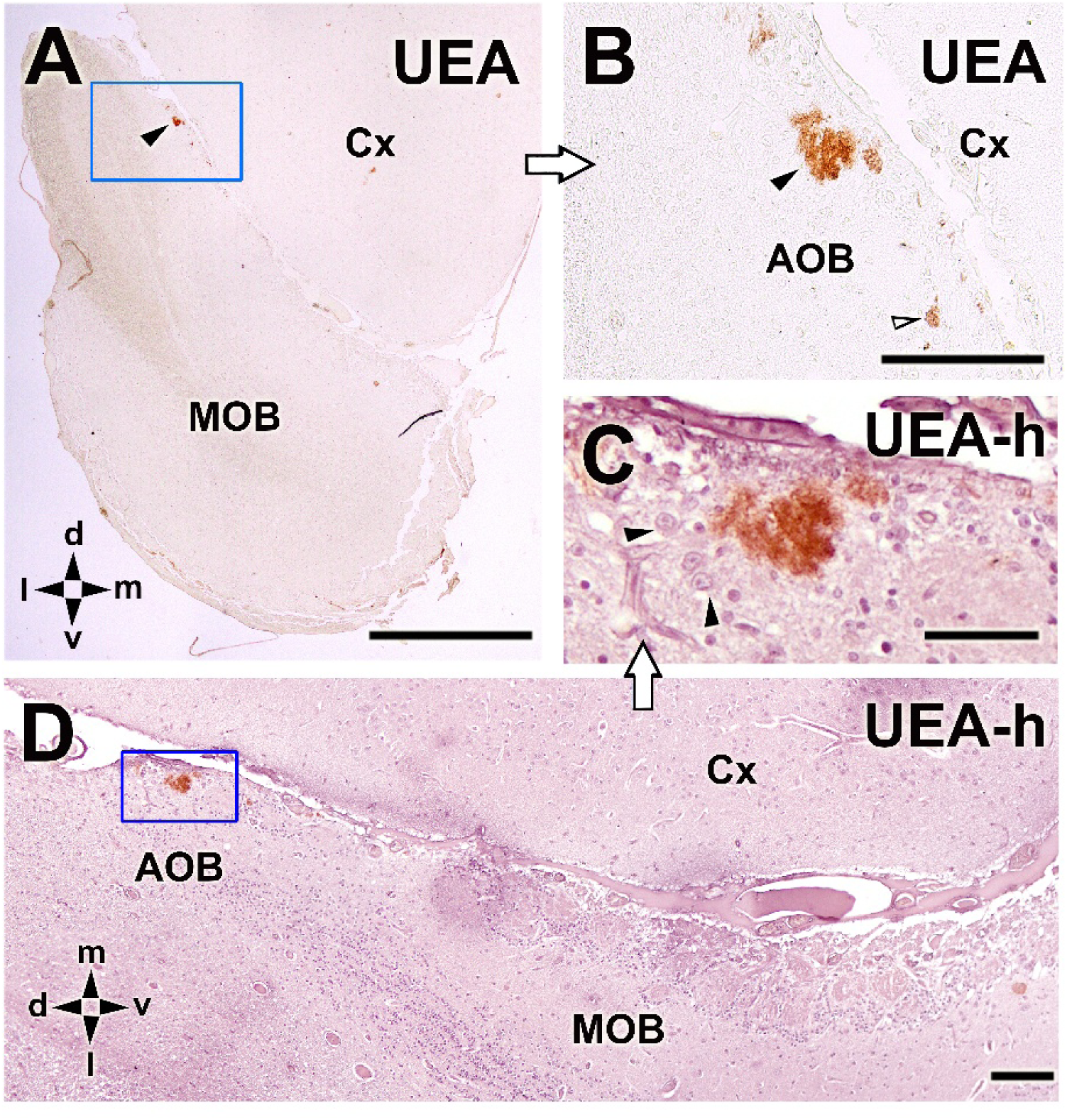
Lectin histochemical analysis of the AOB of *Aonyx cinereus* using *Ulex europaeus* agglutinin (UEA). **(A)** Low-magnification view showing that UEA does not bind to structures within the telencephalon (Cx) or the MOB, except for a restricted region corresponding to the presumptive accessory olfactory bulb (arrowhead inside the box). **(B)** Higher magnification of the UEA-positive area reveals a strongly labeled glomerular-like structure (arrowhead), accompanied by smaller UEA-positive profiles, presumably corresponding to branches of the vomeronasal nerve (open arrowhead). **(C)** Hematoxylin counterstaining (UEA-h) reveals that the UEA-positive structure is surrounded by UEA-negative neurons displaying large, rounded nuclei, together with smaller, also UEA-negative cells distributed throughout the adjacent tissue. **(D)** Low-power view of the AOB and MOB counterstained with hematoxylin demonstrates that UEA labeling is strictly confined to the accessory olfactory area, with no detectable staining in the MOB. Scale bars: A = 1 mm; D= 250 µm; B = 100 µm; C = 50 µm.

LEA lectin histochemistry revealed a distinct labeling pattern in both main and accessory OBs. At low magnification, intense LEA staining was observed in the nerve and glomerular layers of the MOB, as well as in a restricted region corresponding to the presumptive AOB (Fig. 7A). In contrast, the rest of the encephalon showed only diffuse labeling, predominantly associated with perineuronal nets in the cerebral cortex. Higher-magnification analysis of hematoxylin-counterstained sections confirmed the strong LEA positivity of the AOB (Fig. 7B). Detailed examination revealed that, within the MOB, LEA labeling was confined to the nerve and glomerular layers, whereas in the AOB it delineated irregular glomerular-like structures (Fig. 7C,D). Beyond the afferent nerve fibers entering the AOB, no other cellular elements neurons exhibited UEA labeling, including the somata of projection neurons (Fig. 7D). In the cerebral cortex. LEA staining highlighted heterogeneity among neighboring cells, with LEA-positive elements intermingled with unlabeled cells, suggesting differential lectin binding across bulbar subpopulations.

**Figure 7.**
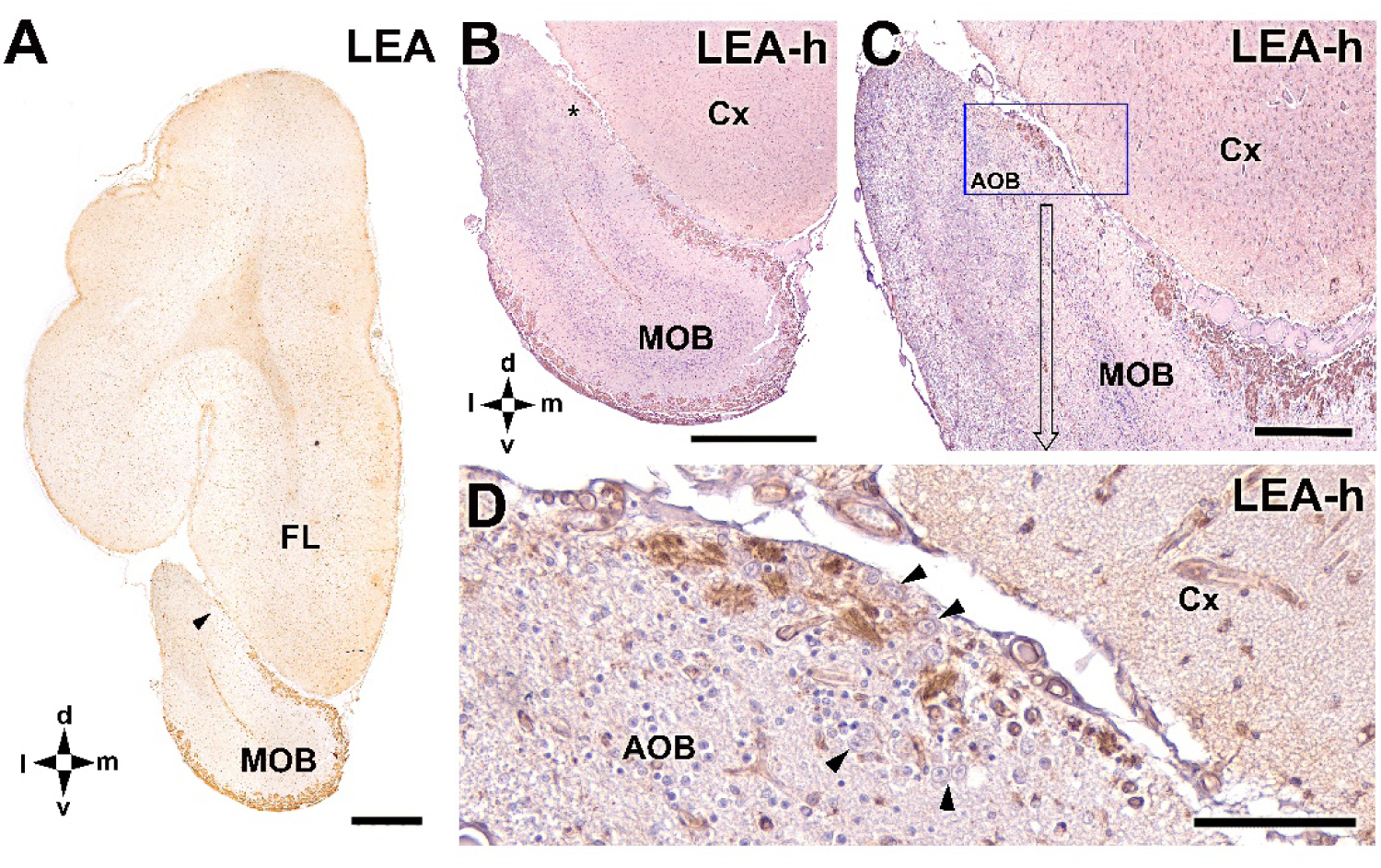
Lectin histochemical analysis of the OB using *Lycopersicon esculentum* agglutinin (LEA). **(A)** Transverse section of the rostral encephalon showing strong LEA labeling in the olfactory nerve, glomerular, and subventricular layers of the MOB, as well as in the presumptive AOB (arrowhead). In the frontal lobe (FL) of the telencephalon, diffuse labeling is observed, mainly associated with perineuronal nets. **(B)** Higher-magnification view of the OB counterstained with hematoxylin (LEA-h), highlighting the LEA-positive region corresponding to the AOB (asterisk) and MOB. **(C,D)** Successive higher-magnification views show that LEA labeling in the MOB is restricted to the nerve and glomerular layers, while in the AOB it appears in irregular glomerular-like structures. Beyond the afferent nerve fibers entering the AOB, no other cellular elements neurons exhibit UEA labeling, including the somata of projection neurons (arrowheads). In the cerebral cortex (Cx), LEA-positive cells coexist with unlabeled cells. Scale bars: A,B = 1 mm; C= 250 µm; D = 100 µm.

STA lectin histochemistry revealed a selective binding pattern within the OB, primarily restricted to superficial layers. In hematoxylin-counterstained sections, STA labeling was observed in the glomerular and nerve layers of the MOB, as well as in the AOB (Fig. 8A). No labeling of deeper cellular layers was detected. Analysis of uncounterstained sections confirmed the specificity of STA for these superficial compartments, clearly delineating the nerve and glomerular layers of the MOB and the corresponding structures of the AOB (Fig. 8B,C). In addition, STA binding was evident in the subventricular zone. Notably, the relative extent and intensity of STA labeling in the AOB were comparable to those observed with UEA and LEA, supporting the consistency of lectin histochemistry as a tool for identifying and labeling accessory olfactory structures.

**Figure 8.**
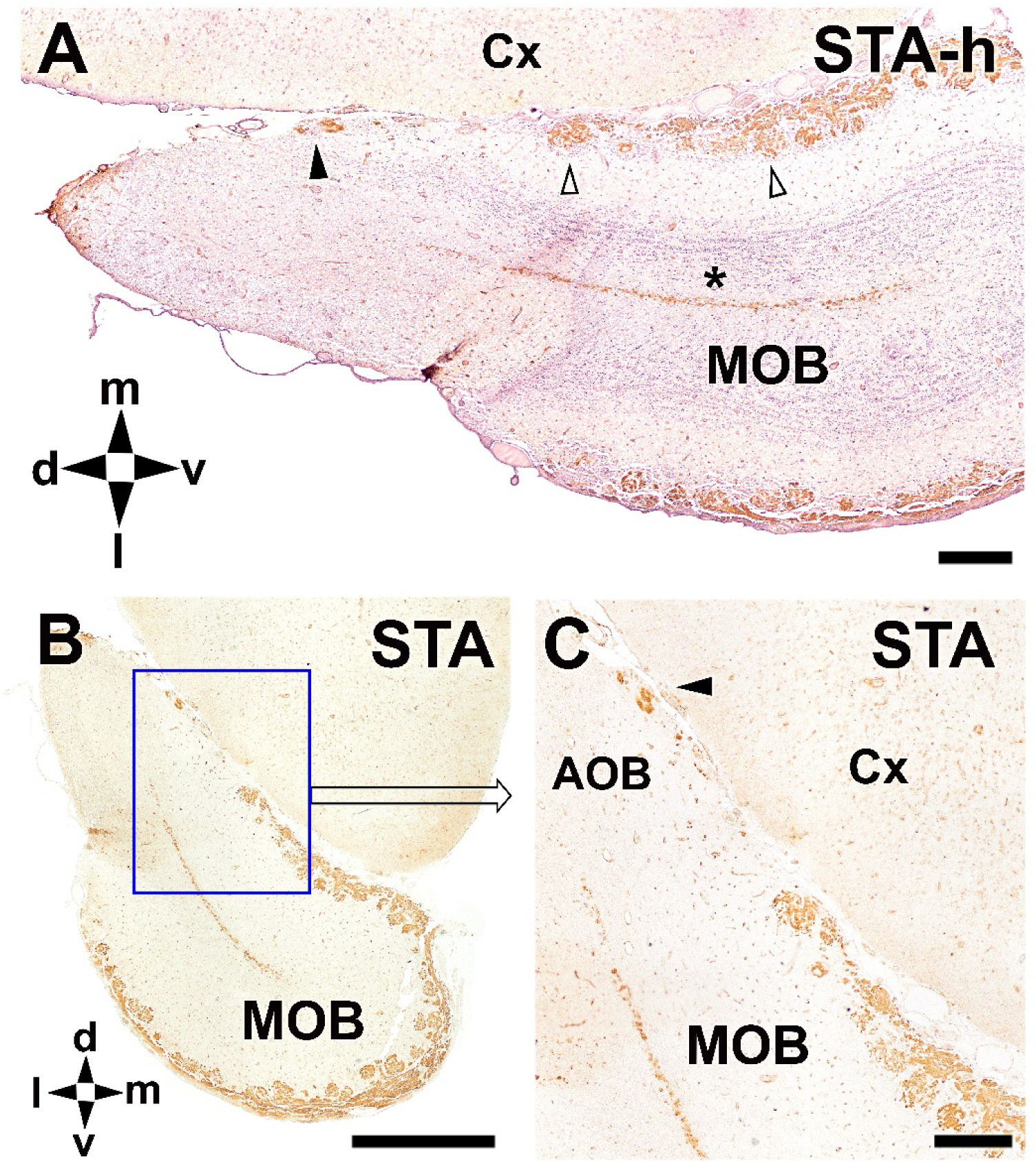
Lectin histochemical analysis of the OB using *Solanum tuberosum* agglutinin (STA). **(A)** Hematoxylin-counterstained section (STA-h) showing STA labeling in the glomerular and nerve layers of both the MOB (open arrowheads) and AOB (arrowhead) and in the subventricular zone (asterisk). Labeling is restricted to superficial layers, with no evident staining of deeper cellular layers. **(B,C)** Uncounterstained sections showing the specificity of STA binding for the superficial layers of the OB, including the nerve and glomerular layers of the MOB and the AOB. The relative extent of STA labeling in the AOB is comparable to that observed with UEA and LEA. Scale bars: B = 1 mm; A,C= 250 µm.

Histochemical analysis using VVA and SBA revealed very low levels of lectin binding in the olfactory bulb. With both lectins, labeling was scarce and restricted to a small number of individual glomeruli located along the peripheral circumference of the MOB (Fig. 9A,C). No widespread staining of olfactory bulb layers or cellular elements was observed. Higher-magnification analysis confirmed that the VVA- and SBA-positive structures corresponded to isolated glomeruli with similar morphology and distribution (Fig. 9B,D). The comparable localization and appearance of the labeled profiles indicate that both lectins recognize in the MOB of the Asian small-clawed otter a limited subset of glomerular structures.

**Figure 9.**
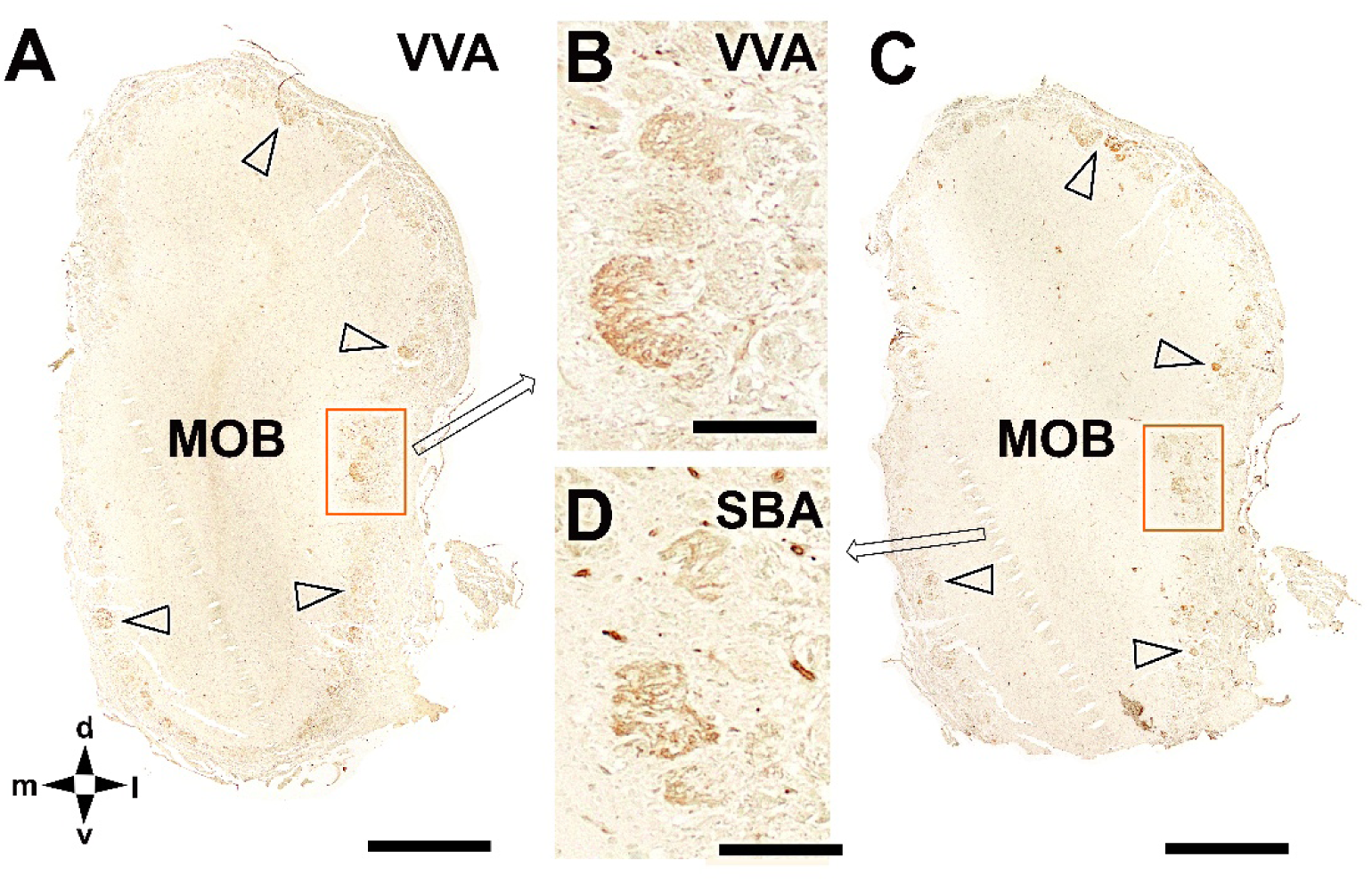
Lectin histochemical analysis of the olfactory bulb using *Vicia villosa* agglutinin (VVA) and *Glycine max* agglutinin (SBA). (**A**) Transverse section of the MOB showing very limited VVA labeling, restricted to a small number of individual glomeruli distributed along the peripheral circumference of the MOB. **(B)** Higher-magnification view of a VVA-positive region illustrating the morphology of the labeled glomeruli. **(C)** Transverse section of the MOB processed with SBA showing a similarly sparse labeling pattern, with a small number of individual glomeruli located around the MOB. **(D)** Higher-magnification view of the SBA-positive glomeruli, revealing a localization and appearance comparable to those observed with VVA. Scale bars: A,C = 1 mm; B,D = 100 µm.

### Immunohistochemical study

Once the presumptive accessory olfactory bulb was identified based on its anatomical location and lectin-binding profile, the next step was to perform an immunohistochemical analysis using a panel of neuronal markers in order to assess the neuronal nature of the elements composing this structure.

Anti-Neural cell adhesion molecule (NCAM) and anti-neuron-specific enolase (NSE) are two well-established markers of neuronal differentiation and maturity. NCAM immunohistochemistry produced a selective labeling pattern within the OB, predominantly affecting superficial layers. At low magnification, NCAM immunoreactivity was detected in the glomerular layer of the MOB and in the presumptive AOB (Fig. 10A). Higher-magnification views confirmed that all glomeruli of the MOB were strongly NCAM-positive (Fig. 10B,C). In the AOB, NCAM labeling encompassed the discrete superficial structures previously identified with lectin histochemistry. In sagittal sections, NCAM-positive profiles were again observed in superficial layers of the MOB, further supporting the association of this marker with primary sensory afferent and glomerular compartments (Fig. 10D).

**Figure 10.**
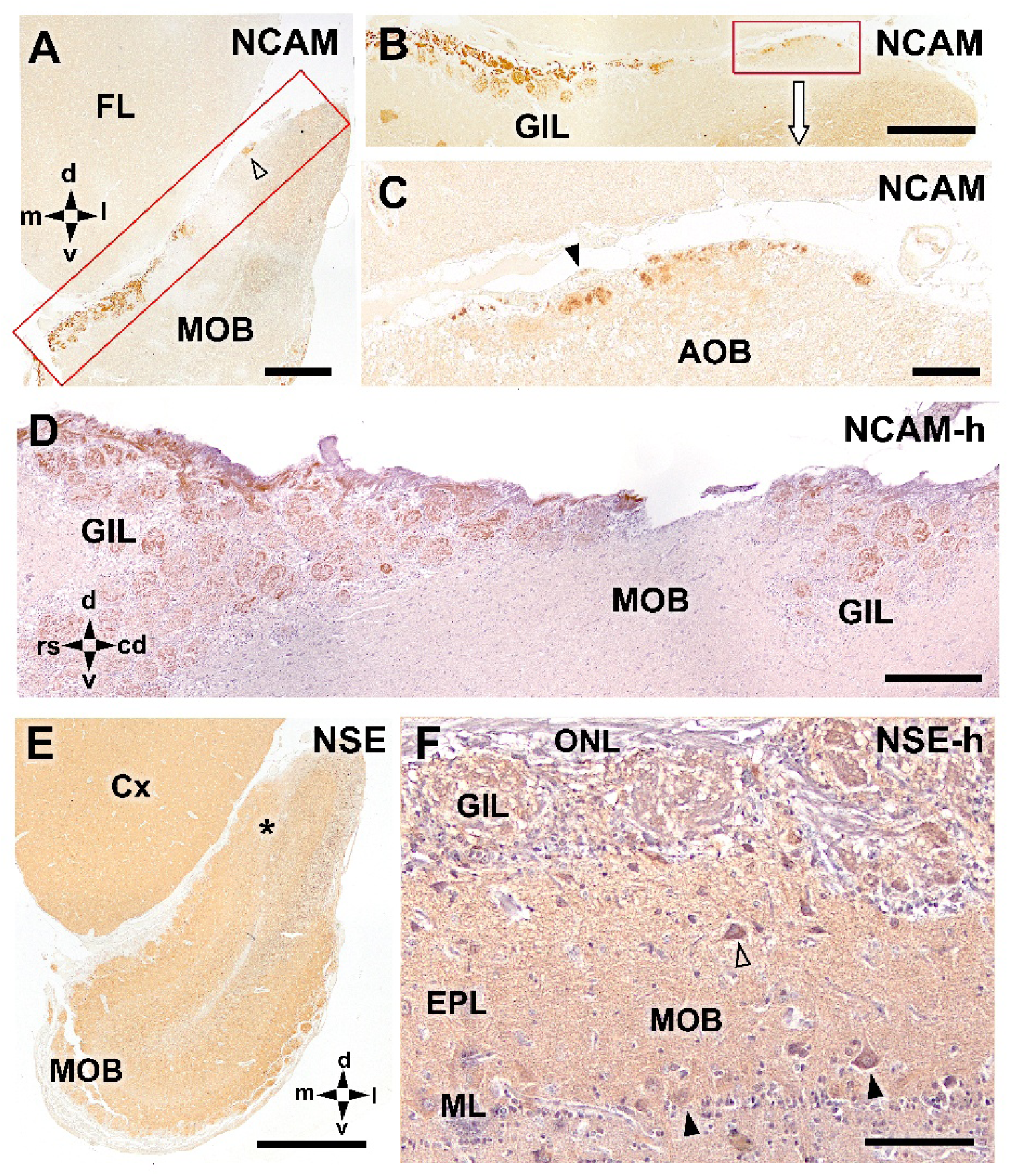
Immunohistochemical analysis of the OB of the Asian small-clawed otter using antibodies against NCAM and neuron-specific enolase (NSE). **(A)** Low-magnification view showing anti-NCAM immunoreactivity in the superficial layers of the MOB and in the presumptive AOB (open arrowhead). **(B,C)** Higher-magnification views of the region indicated in (A) illustrating that NCAM expression is prominent in the superficial layers of the MOB and in the AOB. In the AOB, labeling comprises the neuronal structures previously identified with lectin histochemistry. In the MOB, all glomeruli show anti-NCAM immunoreactivity. **(D)** Sagittal section of the MOB counterstained with hematoxylin shows anti-NCAM-positive structures in the superficial layers, with labeling intensity varying among individual glomeruli. **(E)** Low-magnification view of anti-NSE immunoreactivity demonstrating widespread expression in the frontal lobe (Cx) and the OBs. Within the OB, the superficial layer corresponding to sensory afferents of the olfactory nerve lacks NSE labeling, whereas deeper layers show generalized staining, including in the AOB (asterisk). **(F)** Anti-NSE immunohistochemistry in the MOB, counterstained with hematoxylin, reveals widespread immunoreactivity in both neuronal elements and neuropil. This labelling allows the identification of glomerular structures (GIL), periglomerular cells, tufted cells (open arrowhead), and mitral cells (arrowheads). Scale bars: D,E = 1 mm; A,B= 500 µm; F = 250 μm; C = 100 µm.

Anti-NSE immunohistochemistry showed a markedly different pattern. As expected for a pan-neuronal marker, NSE was broadly expressed throughout the MOB and the frontal lobe (Fig. 10E). Interestingly, within the OB, the superficial nerve layer— corresponding to olfactory nerve afferents—lacked NSE immunoreactivity, whereas deeper layers displayed generalized labeling, including in the AOB (Fig. 10E). At higher magnification of the MOB, NSE immunoreactivity was observed in multiple neuronal populations, including mitral cells, tufted cells, and periglomerular cells in the GlL (Fig. 10F).

Anti-Gαi2 immunohistochemistry revealed a highly specific labeling pattern within the OB. At low magnification, Gαi2 immunoreactivity was confined to a restricted region corresponding to the presumptive AOB, with no detectable labeling in the MOB or in other regions of the encephalon (Fig. 11A). Higher-magnification analysis showed strong immunopositivity within glomerular structures of the AOB (Fig. 11B), consistent with the presence of V1R-associated signaling pathways. In contrast, Gαo immunohistochemistry showed a much broader distribution. At low magnification, intense labeling was observed in the nerve and glomerular layers of the MOB, together with diffuse immunoreactivity throughout the neuropil of both the OB and the surrounding telencephalon (Fig. 11C). Within the AOB, Gαo immunoreactivity was also detected, although with lower intensity compared to Gαi2 (Fig. 11D). These results suggest the presence of functionally differentiated vomeronasal-related signaling components within the presumptive AOB.

**Figure 11.**
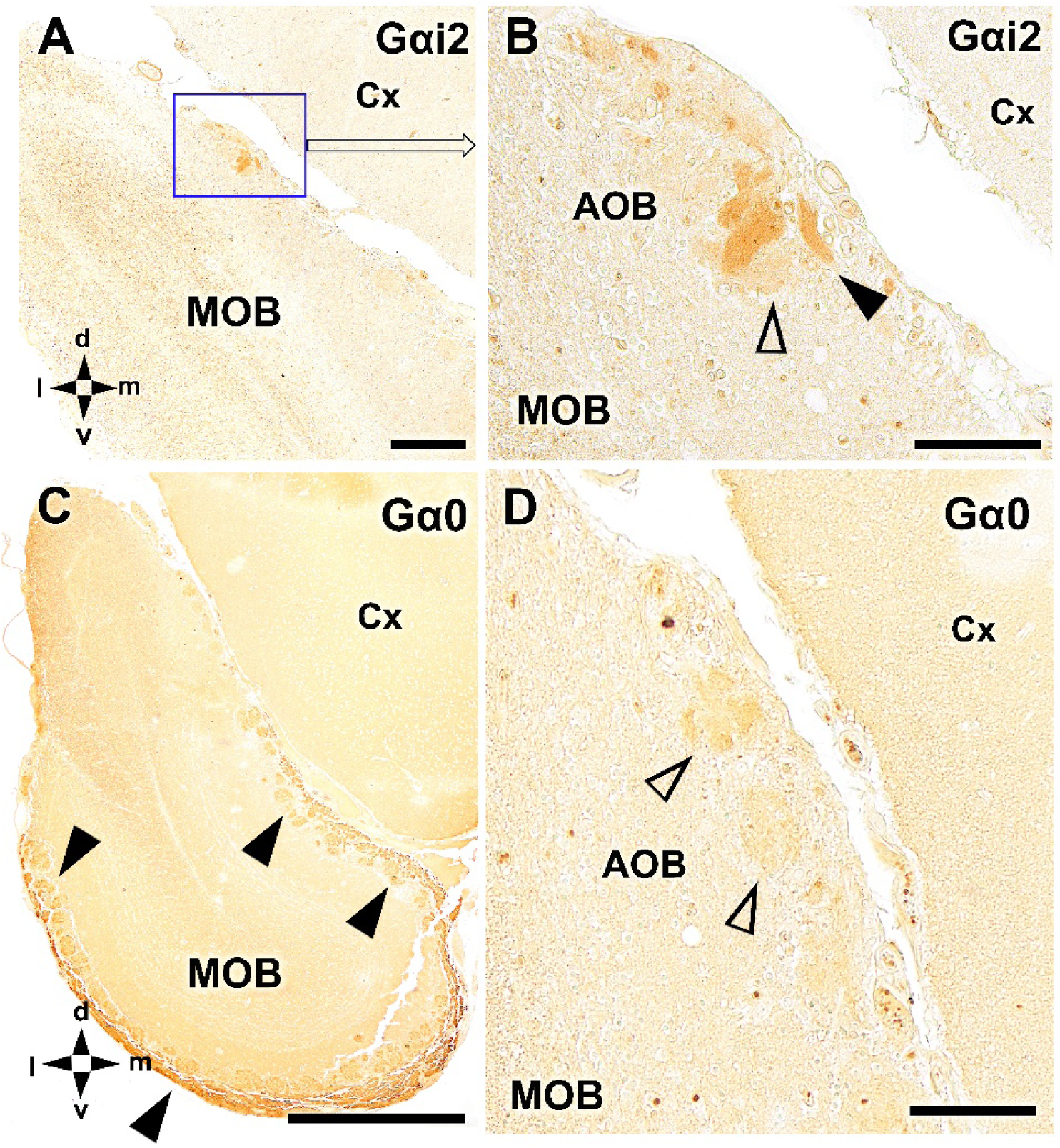
Immunohistochemical study of the olfactory bulb of the Asian small-clawed otter using antibodies against Gαi2 and Gαo. **(A)** Low-magnification transverse section showing anti-Gαi2 immunoreactivity restricted to a discrete region corresponding to the presumptive AOB, with no detectable labeling in the MOB or the surrounding telencephalon. **(B)** Higher-magnification view of the AOB illustrating strong anti-Gαi2 immunopositivity in glomerular structures (arrowheads), but no detectable immunostaining in the deeper layers of the AOB (open arrowhead). **(C)** Low-magnification transverse section processed for anti-Gαo showing intense immunoreactivity in the nerve and glomerular layers of the MOB (arrowheads), as well as diffuse labeling throughout the neuropil of the OB and the telencephalon. **(D)** Higher-magnification view of the AOB area demonstrating weak anti-Gαo immunoreactivity, less intense than that observed for anti-Gαi2. Scale bars: C = 1 mm; A= 250 µm; B,D = 100 µm.

Anti-OMP immunohistochemistry revealed a remarkably difference between the main and accessory OBs. In the MOB, strong anti-OMP immunoreactivity was observed in the superficial layers, particularly within the nerve and glomerular layers, both in transverse and sagittal sections (Fig. 12A,B). However, labeling intensity was heterogeneous, as strongly immunopositive glomeruli were interspersed with weakly labeled or nearly unlabeled ones. In contrast, OMP expression in the presumptive AOB was very limited. In sagittal sections, the AOB was identified caudal to the caudal glomeruli of the MOB (Fig. 12C). In the AOB, anti-OMP immunoreactivity was weak and restricted to a small number of fibers, with no clear labeling of glomerular structures or neuronal cell bodies (Fig. 12D).

**Figure 12.**
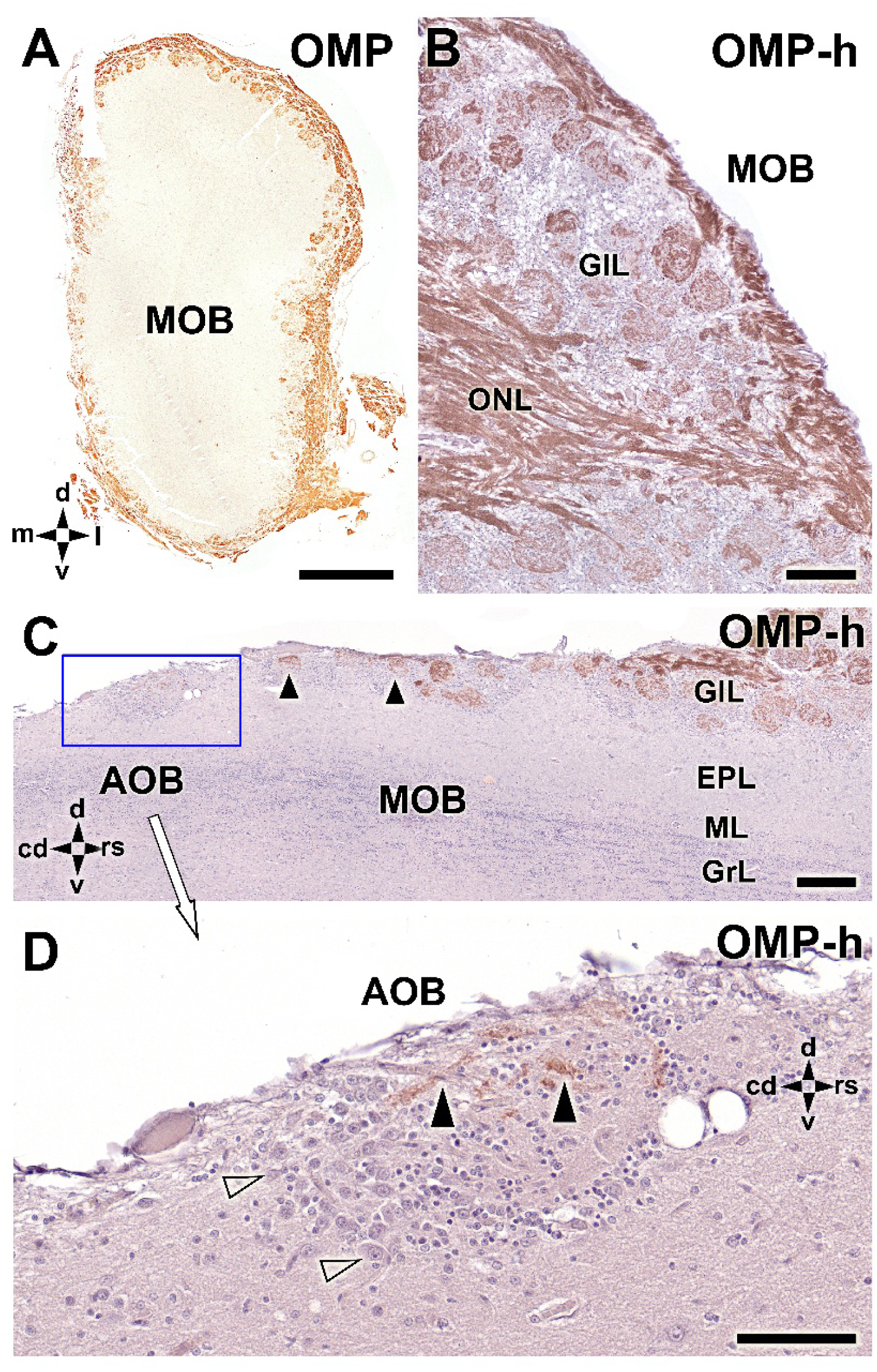
Immunohistochemical analysis of the olfactory bulb of the Asian small-clawed otter using an antibody against olfactory marker protein (OMP). **(A)** Transverse section of the MOB showing strong OMP immunoreactivity in superficial layers. **(B)** Sagittal section of the MOB confirming intense OMP labeling in the glomerular (GlL) and nerve layers (ONL). Not all glomeruli exhibit the same staining intensity, with strongly immunopositive glomeruli intermingled with weakly labeled ones. **(C)** Sagittal section including the caudal region of the MOB and the presumptive AOB. The AOB is located caudal to the caudal most glomeruli of the MOB. **(D)** Higher-magnification view of the AOB showing that OMP labeling is restricted to a small number of fibers, with no evident labeling of glomerular or cellular structures. Scale bars: A = 1 mm; B,C = 250 µm; D = 100 µm.

Immunohistochemical analysis of calcium-binding proteins revealed distinct and complementary labeling patterns in the OB. Anti-CR immunoreactivity was particularly prominent in both the AOB and the glomerular layer of the MOB. In hematoxylin-counterstained sections, the AOB displayed intense anti-CR labeling, while MOB glomeruli showed heterogeneous levels of immunopositivity, with strongly labeled glomeruli interspersed with weakly labeled ones (Fig. 13A). In sections processed without counterstaining, anti-CR immunolabeling was highly specific and highlighted numerous periglomerular cells, as well as neurons located in deeper layers of the OB, including the granular layer (Fig. 13B). Detailed examination revealed distinct patterns of CR expression among different glomeruli, although periglomerular cells were consistently anti-CR-positive regardless of glomerular labeling intensity (Fig. 13D). Higher-magnification analysis of the AOB (Fig. 13C) demonstrated that CR immunoreactivity encompassed both glomerular structures and anti-CR-positive projection neurons, distinguishing them from MOB mitral cells, which lacked CR expression.

**Figure 13.**
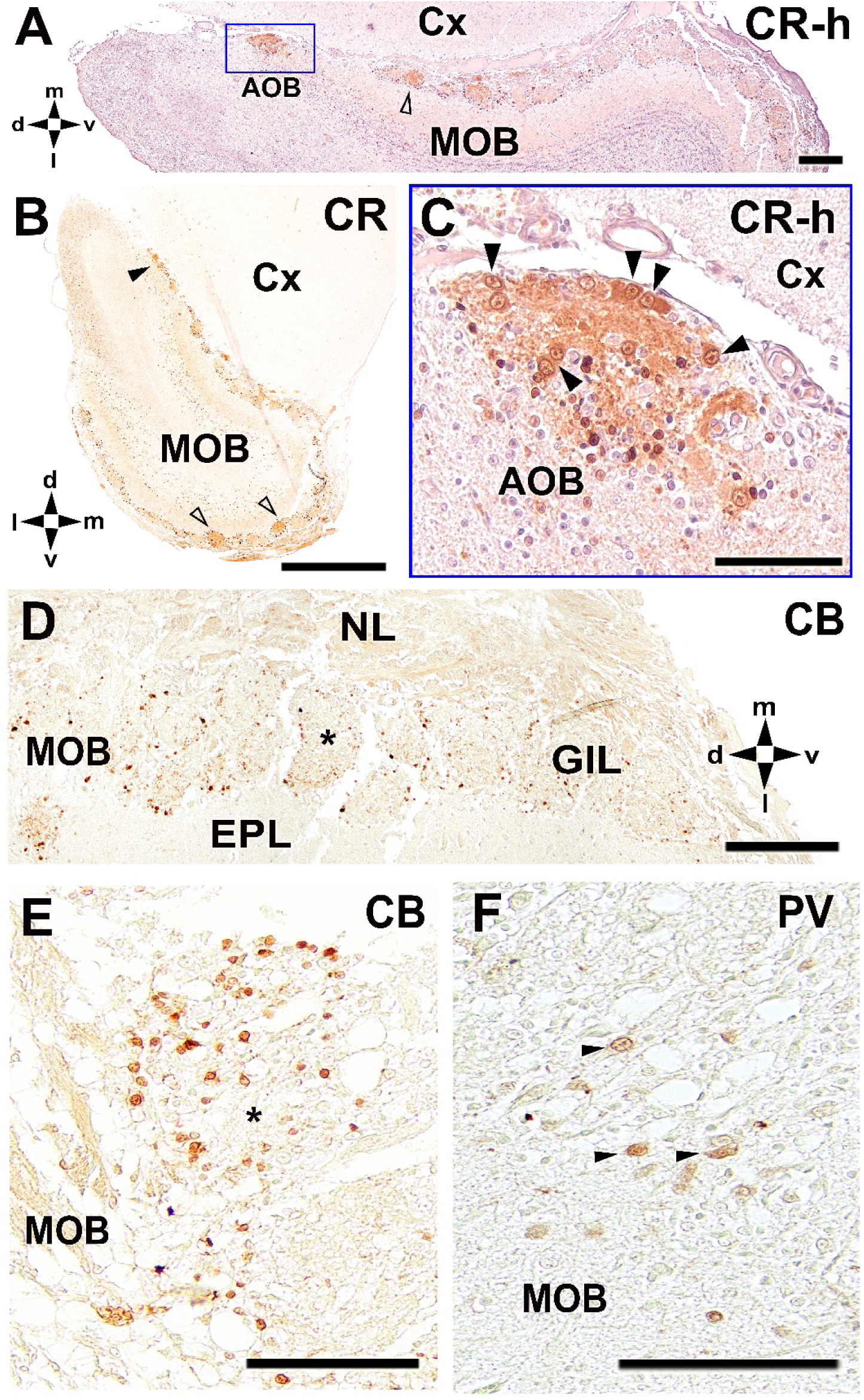
Immunohistochemical analysis of calcium-binding proteins in the olfactory bulb of the Asian small-clawed otter using antibodies against calretinin (CR), calbindin (CB), and parvalbumin (PV). **(A)** Overview of the caudomedial portion of the OB immunostained for anti-calretinin and counterstained with hematoxylin (CR-h). Intense CR immunoreactivity is observed in the AOB and in the GlL of the MOB. Notably, MOB glomeruli display heterogeneous labeling patterns, with some strongly immunopositive and others weakly labeled. **(B)** Low-magnification view of a section processed for anti-CR without counterstaining, showing the overall specificity of CR immunolabeling in the OB. Numerous periglomerular cells are labeled, as well as cells located in deeper layers of the bulb, including the granular layer. The ventral region of the MOB illustrates distinct patterns of CR expression across different glomeruli. Despite this heterogeneity, periglomerular cells are consistently CR-positive across glomeruli (open arrowheads). The AOB region is indicated by an arrowhead. **(C)** Higher-magnification view of the boxed area in (A) showing intense CR immunoreactivity in the AOB, including both glomerular structures and CR-positive projection neurons (arrowheads). **(D,E)** Anti-CB immunohistochemistry reveals labeling restricted to a small subpopulation of neurons, primarily periglomerular cells in the MOB (asterisks). **(F)** Anti-PV immunohistochemistry shows labeling of an even more limited population of periglomerular interneurons in the MOB (arrowheads). Scale bars: B = 1 mm; A,D = 250 µm; C,E, F = 100 µm.

In contrast, calbindin and parvalbumin immunohistochemistry labeled only small and distinct neuronal subpopulations. Anti-Calbindin immunoreactivity was mainly confined to periglomerular cells in the MOB (Fig. 13D,E), whereas parvalbumin labeling was restricted to an even smaller subset of periglomerular interneurons (Fig. 13F).

NeuN immunohistochemistry revealed a generalized neuronal labeling pattern throughout the olfactory system. At low magnification, NeuN immunoreactivity was observed across the MOB, the presumptive AOB, and the cerebral cortex, consistent with its role as a pan-neuronal marker (Fig. 14A). Within both OBs, labeling was particularly pronounced in the glomerular layer. Hematoxylin-counterstained sections further confirmed the strong NeuN immunoreactivity within the AOB (Fig. 14B). At higher magnification, NeuN labeling was evident not only in the neuropil but also in the nuclei of principal neurons within the AOB (Fig. 14C), confirming the neuronal nature of this structure.

**Figure 14.**
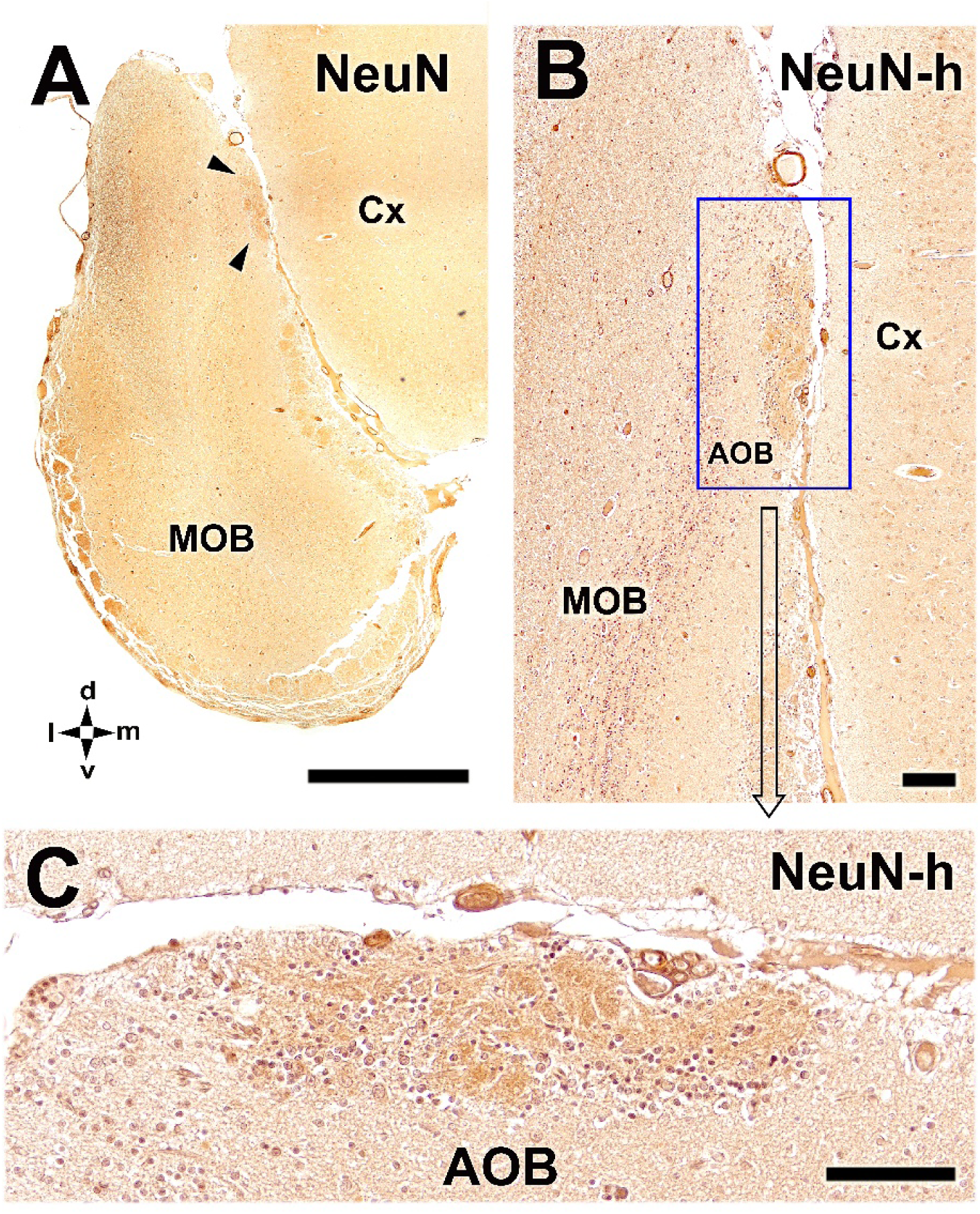
Immunohistochemical analysis of the olfactory bulb of the Asian small-clawed otter using an antibody against NeuN. **(A)** Low-magnification transverse section showing widespread NeuN immunoreactivity throughout the olfactory system, including the MOB, and the presumptive AOB (arrowheads), and the cerebral cortex (Cx). In both OBs, NeuN labeling is particularly intense in the glomerular layer. **(B)** Hematoxylin-counterstained section (NeuN-h) showing the intense anti-NeuN immunoreactivity in the AOB region. **(C)** Higher-magnification view of the AOB shown in (B), demonstrating strong NeuN labeling of both the neuropil and the nuclei of principal neurons. Scale bars: A = 1 mm; B = 250 µm; C = 100 µm.

PGP 9.5 immunohistochemistry revealed a broad neuronal labeling pattern throughout the OB. In sagittal sections of the main MOB, PGP immunoreactivity showed both neuronal somata and sensory afferent fibers entering via the olfactory nerve (Fig. 15A–C). The diversity of neuronal morphologies within the MOB was readily apparent, including distinct populations of mitral cells and tufted cells, as well as periglomerular neurons within the glomerular layer. In a transverse section of the AOB, PGP labeling was more homogeneous and predominantly neuropil-associated (Fig. 15D). Individual neuronal somata were less prominently labeled than with other neuronal markers, resulting in a more diffuse appearance of the AOB. However, analysis of sagittal sections of the AOB revealed numerous principal neurons and several irregularly shaped glomerular structures delineated by surrounding periglomerular cells (Fig. 15E).

**Figure 15.**
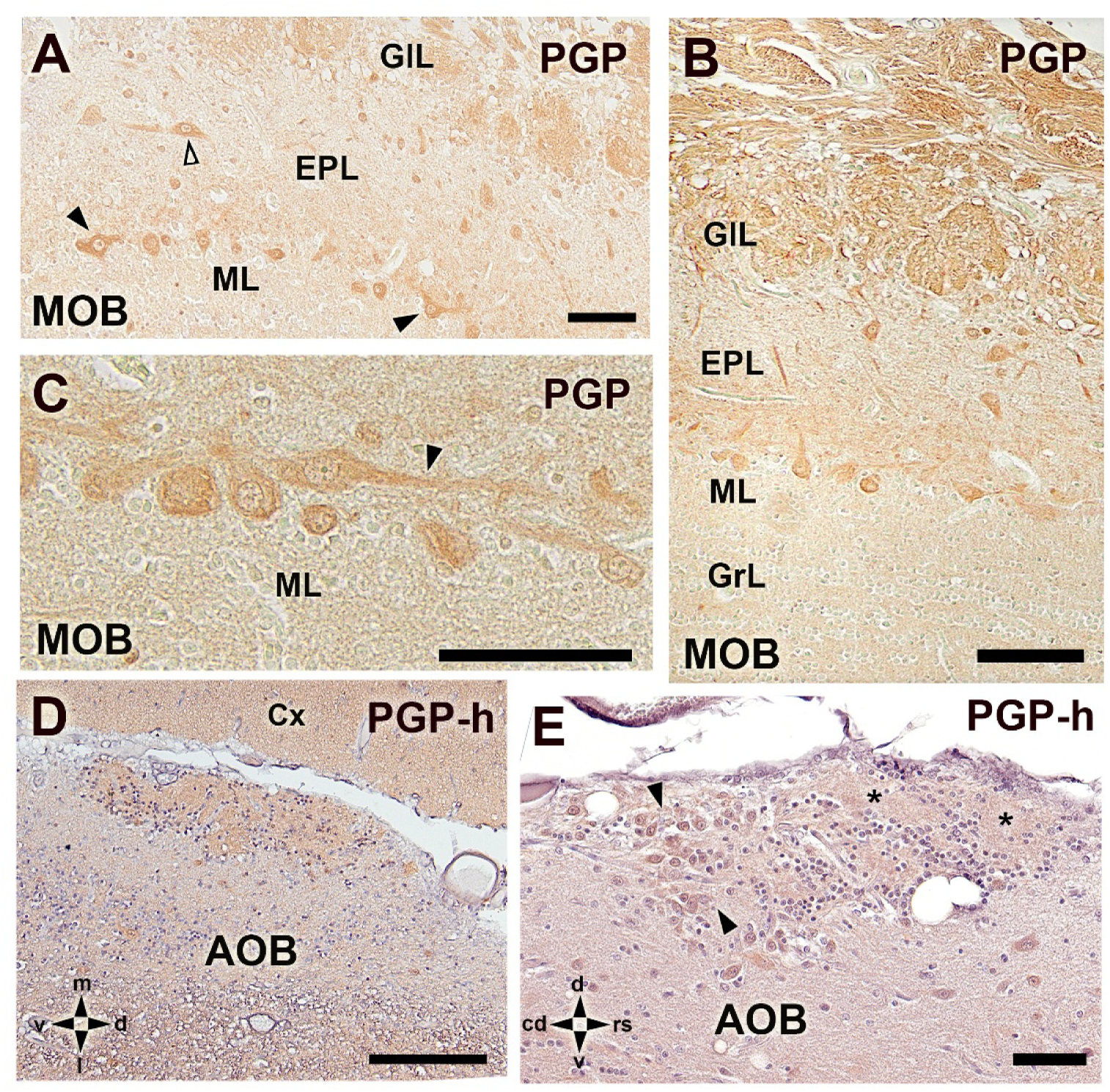
Immunohistochemical analysis of the olfactory bulb of the Asian small-clawed otter using an antibody against protein gene product 9.5 (PGP). (A–C) Sagittal sections of the olfactory bulb showing widespread PGP immunoreactivity in neuronal elements of the MOB, as well as in sensory afferents arriving via the olfactory nerve (B). Distinct morphological types of projection neurons are evident, including mitral cells (black arrowheads) and tufted cells (open arrowheads), together with periglomerular cells within the glomerular layer (GIL). **(D)** Transverse section of the AOB showing generalized PGP labeling of the neuropil. **(E)** Sagittal section of the AOB showing abundant principal cells (arrowheads) and several ill-defined glomeruli (asterisks) delineated by periglomerular cells. Scale bars: A-E = 100 µm.

Anti-tubulin immunohistochemistry provided a clear delineation of OB layers based on their cytoskeletal organization. In the MOB, tubulin labeling revealed a marked compartmentalization, with the GrL showing very low immunoreactivity and the EPL layer displaying intense labeling (Fig. 16A). At higher magnification, the contrast between these layers was particularly evident. The mitral cell layer was readily identifiable, with tubulin-positive dendritic processes extending toward the EPL and axonal profiles projecting toward deeper regions (Fig. 16B). Analysis of transverse sections including the AOB demonstrated that the AOB exhibits a tubulin immunoreactivity pattern similar to that observed in the superficial layers of the MOB (Fig. 16C). Higher-magnification views confirmed that both the MOB and the AOB contain tubulin-positive cellular elements and neuropil, indicating comparable cytoskeletal organization (Fig. 16D,E).

**Figure 16.**
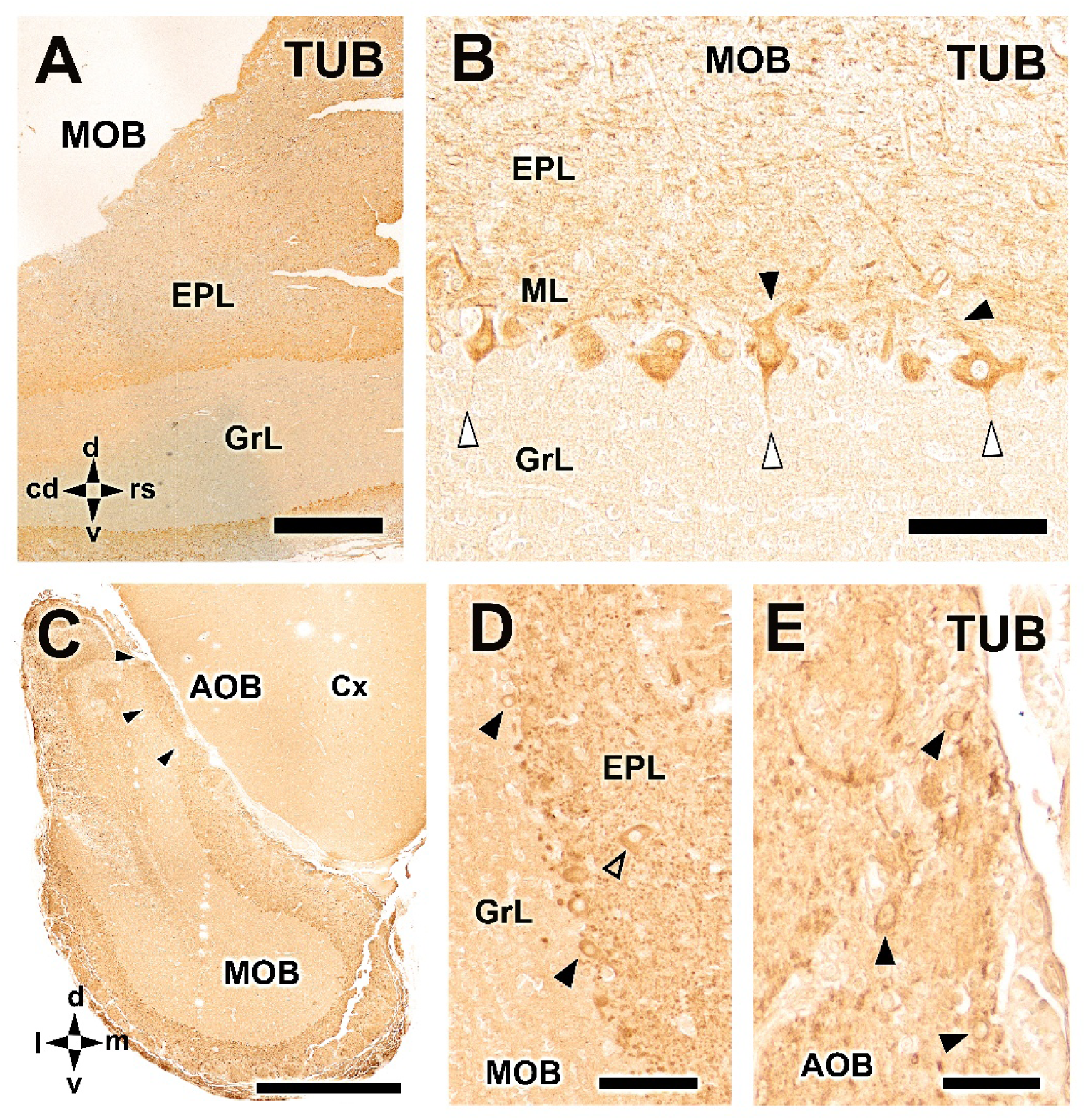
Immunohistochemical analysis of the OB of the Asian small-clawed otter using an antibody against tubulin (TUB). **(A)** Low-magnification view of the MOB showing a clear compartmentalization of layers based on tubulin immunoreactivity. The GrL displays very weak labeling, whereas the EPL is intensely immunopositive. **(B)** Higher-magnification view showing the sharp contrast between the weakly labeled GrL and the strongly labeled EPL. The mitral cells are clearly identifiable, with tubulin-positive dendritic processes (arrowheads) extending toward the EPL and axonal profiles (open arrowheads) projecting toward deeper layers. **(C)** Low-magnification transverse section of the olfactory peduncle including the AOB and the MOB, showing that the AOB exhibits a tubulin immunoreactivity pattern comparable to that observed in the superficial layers of the MOB. **(D)** Higher-magnification view of the MOB in (C) illustrating anti-tubulin immunopositivity in the mitral (arrowheads) and tufted cells (open arrowhead), and in the EPL neuropil. **(E)** Higher-magnification view of the AOB in (C) showing strong anti-tubulin immunoreactivity in the principal cells. Scale bars: A,C = 1 mm; B,D = 100 µm; E = 50 µm.

## DISCUSSION

The present study provides the first detailed anatomical, histological, lectin-histochemical, and immunohistochemical characterization of the main and accessory olfactory bulbs of a lutrine species. The most striking finding is the marked contrast between a prominent and canonically laminated main olfactory bulb (MOB) and a small, macroscopically indistinguishable accessory olfactory bulb (AOB) displaying a highly unconventional cytoarchitecture. Despite its limited size and the absence of clear lamination, the identity of the AOB was consistently supported by its topographical location, neuronal composition, selective lectin-binding profile, and strong Gαi2 immunoreactivity. These findings demonstrate that the accessory olfactory pathway is retained in *Aonyx cinereus*, although its first central relay exhibits a structurally reduced but neurochemically differentiated organization.

Mammals occupy an exceptionally broad range of ecological niches and exhibit considerable diversity in their social organization, reproductive strategies, territorial behaviour, and reliance on different sensory modalities. Chemical cues contribute to many of these processes, including individual and sexual recognition, assessment of reproductive status, establishment of social hierarchies, territorial marking, maternal behaviour, predator avoidance, and prey detection (Nolte et al. 1994; Kimchi et al. 2007; Khallaf et al. 2026; Numan 2026; Yang et al. 2026). The main and accessory olfactory systems should therefore not be regarded as anatomically invariant pathways but as dynamic sensory systems whose organization may reflect the interaction between their phylogeny, ecology, and behaviour. Most current knowledge concerning the mammalian vomeronasal system derives from laboratory rodents, particularly mice and rats. These species exhibit a well-defined VNO, a clear apical-basal subdivision of their sensory epithelium, segregated Gαi2- and Gαo-associated projections, and a laminated AOB divided into anterior and posterior domains (Vaccarezza et al. 1981; Tirindelli and Ryba 1996; Salazar and Sanchez-Quinteiro 1998; Vargas-Barroso et al. 2017; Kondoh et al. 2018). Comparative studies, however, have demonstrated substantial interspecific variation in sensory epithelial differentiation, receptor repertoires, expression of transduction proteins, projection patterns, and AOB cytoarchitecture (Takigami et al. 2000; Takigami 2004; Kondoh et al. 2022; Torres et al. 2023a; Sanmartín-Vázquez et al. 2024; Shin et al. 2026). Consequently, the rodent condition cannot be assumed to represent a universal mammalian pattern. This comparative framework is also relevant to species showing regression of canonical vomeronasal components, including aquatic mammals and humans, because it helps distinguish conserved structures from lineage-specific specializations and secondary reductions without implying direct functional equivalence between distantly related taxa (Switzer, III. et al. 1980; Smith et al. 2001; Hecker et al. 2019; Kondoh et al. 2024).

### The accessory olfactory system in carnivorans and mustelids

Carnivorans are particularly informative for this comparative approach because they encompass species with markedly different ecological and social characteristics. Although chemical communication is important throughout the order, the organization of the vomeronasal system differs considerably among families and even between closely related species (Tomiyasu et al. 2017; Torres et al. 2021; Itakura et al. 2026). Canids provide a clear example. The domestic dog possesses a relatively small and poorly differentiated AOB (Miodonski 1968; Nakajima et al. 1998; Salazar et al. 2013), whereas studies in wild canids, including the red fox and wolf, have revealed a more readily identifiable and neurochemically complex accessory pathway (Chengetanai et al. 2020; Ortiz-Leal et al. 2023; Ortiz-Leal et al. 2024). Thus, the absence of a conspicuous bulbar prominence or conventional lamination does not necessarily imply loss of the AOB.

The same interpretative problem is particularly evident in Mustelidae. Members of this family occupy terrestrial, fossorial, arboreal, freshwater, and marine environments and display considerable variation in their social and reproductive behaviour. Scent marking is widespread among mustelids and frequently involves urine, faeces, and secretions from specialized glands that may convey information about identity, territorial occupancy, reproductive condition, and social status (Hutchings and White 2000; Noonan et al. 2019). Nevertheless, the neural substrates involved in processing these signals remain poorly characterized.

The few available studies indicate that mustelids do not necessarily conform to the classical rodent organization. In the mink, *Neogale vison* -formerly *Mustela vison*- the VNO contains a differentiated sensory epithelium and the characteristic vascular and structural components of a peripheral vomeronasal organ (Salazar et al. 1994a; Salazar et al. 1997). A presumptive AOB was identified using morphology and lectin histochemistry, although its organization differed from that described in commonly studied rodents (Salazar et al. 1998b). In the domestic ferret, the VNO is also present, whereas the AOB is relatively small, poorly laminated, and difficult to delimit using conventional histological criteria (Weiler 1999; Kelliher et al. 2001; Woodley et al. 2004). These features probably contributed to earlier descriptions of the AOB of some mustelids as absent, rudimentary, or poorly defined (Jawlowski 1956). The organization observed in *A. cinereus* therefore appears to fit a broader mustelid tendency in which the AOB is retained but cannot be reliably recognized according to the canonical rodent model.

### Lutrines and the secondary transition to aquatic environments

The interpretation of olfactory structures in Lutrinae must also consider their secondary adaptation to aquatic and semiaquatic environments. This transition modifies the physical conditions under which chemical information is produced, dispersed, and detected. Volatile compounds behave differently in air and water, whereas non-volatile substances deposited in secretions, urine, or faeces may persist on terrestrial substrates bordering aquatic habitats (Catania 2006; Mollo et al. 2014; Mollo et al. 2017). Consequently, an aquatic or semiaquatic lifestyle need not produce uniform regression of the olfactory systems; instead, it may favour differential retention or modification of particular sensory components.

Comparisons among aquatic mammals support this heterogeneous evolutionary pattern. Odontocetes have lost recognizable olfactory bulbs and associated central pathways, whereas mysticetes retain a reduced main olfactory system and a limited repertoire of functional olfactory receptor genes (Kishida et al. 2015b; Kishida et al. 2015a; De Vreese et al. 2023). The VNO and its associated accessory pathways are generally considered absent in cetaceans (Thewissen et al. 2011; Berta et al. 2014). Pinnipeds display a more variable condition: no sensory VNO or recognizable AOB was identified in the harbour seal, whereas a differentiated VNO and a V1R–Gαi2-associated pathway have recently been described in the Steller sea lion (Kondoh et al. 2024; Kondoh et al. 2025a). Similarly, differentiated vomeronasal structures in the capybara and Eurasian beaver demonstrate that semiaquatic adaptation does not necessarily implies loss of the VNS (Suárez et al. 2011; Tomiyasu et al. 2022).

### Morphological retention and reduction of the vomeronasal system in Lutrinae

Recent evidence from *Lontra canadensis* provides an important peripheral counterpart to the central organization described here. Using iodine-enhanced computed tomography and comparisons with the American mink (*Neogale vison*) and long-tailed weasel *(Neogale frenata*), Palmer et al. (2026) confirmed the persistence of the VNO in a semiaquatic lutrine. However, the river otter VNO differed from the other mustelids examined in several respects, including a less expanded lumen and a comparatively large space between the organ and its surrounding cartilaginous capsule. These features were interpreted as potentially consistent with a degree of structural reduction. Moreover, vascular structures comparable to those observed around the mink VNO **(Salazar et al. 1994a)** were not clearly resolved in the river otter using diceCT, although the absence of contrast-enhanced vascular profiles cannot be considered equivalent to histological absence. Collectively, these observations suggest that retention of the VNO in lutrines may be accompanied by substantial modification of its internal organization and supporting structures. The persistence of vomeronasal structures despite molecular regression may also reflect the developmental integration of the VNO with surrounding nasal and palatal tissues. Formation of the vomeronasal cartilage, incisive duct, nasal septum, and adjacent palatal structures involves interacting morphogenetic environments that may allow some components of the vomeronasal complex to persist even when selective pressure on sensory receptor repertoires has been relaxed (Salazar et al. 2003; Smith et al. 2014; Palmer et al. 2026). Such developmental integration offers a plausible explanation for the anatomical persistence of peripheral structures, although it does not by itself account for the preservation of a neurochemically differentiated central relay such as the AOB identified in *A. cinereus*.

The present study complements those peripheral observations by demonstrating preservation of the first central vomeronasal relay in another lutrine. Whereas Palmer et al. (2026) identified the VNO in *L. canadensis*, our findings reveal a neurochemically differentiated AOB in *A. cinereus*. Together, these studies indicate that the transition towards semiaquatic life has not resulted in complete disappearance of the vomeronasal pathway in Lutrinae. Instead, the system appears to have undergone lineage-specific remodelling involving unequal preservation of its peripheral, molecular, and central components. This interpretation is consistent with the ecology of *A. cinereus*. Asian small-clawed otters inhabit rivers, wetlands, and rice fields and repeatedly use latrine sites at which chemical deposits can persist and be encountered by conspecifics (Hussain et al. 2011; Aadrean and Usio 2017). Some latrine sites remain active for several years, supporting their significance as stable communication points within semiaquatic landscapes (Aadrean and Usio 2017). Retention of both main and accessory olfactory structures may therefore be relevant to a species that combines aquatic foraging with extensive social interactions and chemical communication on terrestrial substrates.

### Macroscopic and microscopic organization of the MOB

The prominent and elongated morphology of the olfactory bulbs, together with the development of the olfactory peduncles, olfactory tubercle, piriform lobe, and lateral olfactory tract, indicates substantial preservation of the main olfactory pathway in *A. cinereus*. This finding is notable in the context of semiaquatic adaptation because it differs markedly from the regression or loss of olfactory structures reported in several fully aquatic mammalian lineages (De Vreese et al. 2023). The macroscopic appearance of the MOB is therefore consistent with a continued biological relevance of main olfaction in lutrines.

Histologically, the MOB exhibited the canonical mammalian lamination, comprising olfactory nerve, glomerular, external plexiform, mitral cell, internal plexiform, and granular layers. This conserved arrangement indicates that the principal stages of bulbar information processing remain anatomically segregated, as occurs in other mammals with a well-developed main olfactory pathway (Firestein 2001; Ashwell 2006; Tufo et al. 2022). The abundance and morphological diversity of mitral and tufted cells, together with well-defined populations of periglomerular and granule cells, further indicate a complex organization of projection neurons and local interneuronal circuits. The dense and clearly delineated lateral olfactory tract visualized using Tolivia staining provides additional evidence for the development of MOB output pathways. Mitral and tufted cell axons converge within this tract before projecting towards primary olfactory cortical regions, including the olfactory tubercle, piriform cortex, and amygdaloid areas (Switzer, III. et al. 1980; Schwob and Price 1984; Inaki et al. 2004; Baum and Cherry 2015). Its prominent myelination in *A. cinereus* is therefore consistent with the anatomical differentiation observed throughout the MOB.

### A small and non-canonically organized AOB

In contrast to the MOB, no structure attributable to the AOB could be identified by external inspection. Serial transverse and sagittal histological sections revealed only a restricted dorsocaudal region within the olfactory peduncle, located immediately caudal to the most posterior MOB glomeruli. This position corresponds to the expected location of the mammalian AOB, although the structure lacked the clear surface prominence observed in rodents, lagomorphs, ruminants, and marsupials (Schneider et al. 2012; Villamayor et al. 2020; Kondoh et al. 2025b; Fernandez-Aburto et al. 2026). The cytoarchitecture of this region differed markedly from the canonical laminated AOB. A continuous glomerular layer, a clearly demarcated mitral-plexiform layer, and a distinct granule cell layer could not be recognized. Instead, the presumptive AOB contained a limited number of irregular glomerular-like structures and a heterogeneous mixture of neuropil, large neurons with prominent nuclei and nucleoli, and smaller cells resembling granule neurons. The larger cells probably correspond to AOB projection neurons, but their arrangement did not form the continuous mitral-plexiform layer characteristic of the rodent AOB (Salazar et al. 2006; Larriva-Sahd 2008).

This pattern resembles the poorly differentiated organization reported in the domestic ferret and, to some extent, the mink (Salazar et al. 1998b; Kelliher et al. 2001). It suggests that the mustelid AOB may be organized as a compact and irregular neuronal aggregation rather than as a sharply laminated cortical structure. Accordingly, failure to identify canonical layers should not be interpreted as evidence that the AOB is absent. In mustelids, its recognition appears to require integration of topographical, cellular, histochemical, and molecular criteria.

### Lectin-histochemical characterization

Lectins are particularly valuable for studying olfactory pathways because they recognize glycoconjugates associated with specific sensory neuronal populations, axonal projections, and glomerular compartments. Their binding patterns vary between species, reflecting differences in glycosylation and neuronal organization, but selected lectins can provide informative markers of the main and accessory systems (Plendl and Sinowatz 1998; Salazar et al. 2001; Shin et al. 2017). Among the lectins examined, UEA provided the clearest histochemical evidence for the identity of the AOB. Its binding was restricted to a small region corresponding precisely to the structure identified topographically and cytoarchitecturally as the presumptive AOB. Within this region, UEA strongly labeled a glomerular-like structure and smaller profiles compatible with branches of the vomeronasal nerve, whereas no equivalent labeling occurred in the MOB or adjacent telencephalon. This selective pattern differentiates the accessory region from the glomeruli of the MOB and supports its association with a molecularly distinct sensory projection. Comparable selectivity of UEA-I for components of the vomeronasal pathway has been reported in several mammals. In the mouse, UEA-I predominantly labels Gαi2-expressing apical vomeronasal sensory neurons, their axons, and their glomerular targets in the anterior AOB (Barrios et al. 2014; Kondoh et al. 2017). In canids, UEA-I has also proved particularly useful for distinguishing the accessory pathway: it labels the vomeronasal nerves and superficial layers of the AOB in the domestic dog, red fox, and Iberian wolf, whereas the MOB is unreactive (Salazar et al. 1994b; Ortiz-Leal et al. 2022b; Ortiz-Leal et al. 2024). The highly restricted labeling observed in *A. cinereus* therefore resembles the pattern described in these canids and supports the identification of the UEA-positive region as the AOB. Remarkably, however, this selectivity is not shared by all mustelids. In the American mink, UEA-I was the only one of the four lectins examined that did not bind exclusively and consistently to the presumptive AOB; this region was instead selectively delineated by SBA, BSI-B₄, and VVA (Salazar et al. 1998b). UEA-I should therefore be regarded as a species-dependent histochemical marker of vomeronasal projections rather than as a universal marker of the accessory olfactory pathway.

LEA and STA produced a broader pattern in *A. cinereus*, labeling superficial afferent compartments in both the MOB and AOB. Within the MOB, their binding was largely confined to the nerve and glomerular layers, whereas in the accessory region they delineated irregular glomerular-like structures. Both lectins recognize glycoconjugates containing N-acetylglucosamine residues, and comparable labeling of sensory axons and glomerular structures in both the MOB and AOB has been consistently reported across the mammalian species examined to date (Taniguchi et al. 1993b; Halpern et al. 1998; Torres et al. 2021; Keller et al. 2022; Ortiz-Leal et al. 2024). Their distribution in *A. cinereus* suggests that the superficial compartments of both bulbs retain related glycoconjugate profiles despite their strikingly different cytoarchitectures.

VVA and SBA, by contrast, recognized in *A. cinereus* only a small number of individual glomeruli distributed around the MOB. Interestingly, Salazar et al. (1998b) described a more complex organization of the AOB in the American mink, consisting of a main dorsocaudal eminence and two additional, smaller lateral and medial regions, identified through lectin histochemistry and morphological examination. These latter regions, located in areas generally associated with the MOB, were interpreted as additional components of the accessory olfactory system. However, their precise anatomical identity warrants further consideration, particularly in light of the heterogeneous glomerular populations now recognized within the MOB. In contrast, no reactivity to either lectin was detected in the AOB of

*A. cinereus* in the present study.

Comparable lectin-defined glomerular subsets have been described in other species, although the identity and distribution of the labeled populations vary substantially according to taxon and histochemical protocol (Takami et al. 1992; Plendl and Sinowatz 1998; Tisay et al. 2002). These observations support a chemically heterogeneous organization of the MOB but do not by themselves identify the odorant receptors or functional categories represented by the labeled glomeruli.

### Immunohistochemical characterization of the accessory and main olfactory bulbs

Immunohistochemistry provided several independent lines of evidence supporting the identification and neurochemical characterization of the AOB. The most relevant result was the highly restricted expression of Gαi2, which was confined to glomerular-like structures within the accessory region and was not detected in the surrounding MOB or telencephalon. In the canonical rodent and marsupial vomeronasal pathway, Gαi2 is associated with apical V1R-expressing sensory neurons whose axons project predominantly to the anterior AOB (Halpern et al. 1995; Halpern et al. 1998; Takigami 2004). Comparable Gαi2-positive projections have been described in several other mammalian groups, although their relative development and spatial segregation vary considerably among species (Schneider et al. 2012; Torres et al. 2022; Kondoh et al. 2025b; Ruiz-Rubio et al. 2026).

Gαo immunoreactivity showed a markedly different distribution. Within the AOB, labeling was weak and did not define a clearly segregated glomerular domain comparable to the posterior Gαo-positive subdivision of the rodent AOB. This pattern may indicate a comparatively reduced representation of the canonical V2R-Gαo pathway, spatial intermingling of different vomeronasal projections, or a lutrine-specific organization lacking the clear anterior-posterior compartmentalization described in rodents. Nevertheless, weak immunoreactivity cannot be interpreted as evidence for the complete absence of V2R-associated neurons, particularly because Gαo expression may vary among species and its widespread distribution in the central nervous system reduces its specificity as a vomeronasal marker (Ortiz-Leal et al. 2024). This latter point was illustrated by the MOB, where Gαo immunoreactivity was considerably stronger than in the AOB. Labeling was particularly intense in the olfactory nerve and glomerular layers and was accompanied by diffuse immunoreactivity throughout the olfactory and telencephalic neuropil. A broad distribution of Gαo has also been described in the mouse olfactory system, where its expression is not restricted to vomeronasal sensory projections (Wekesa and Anholt 1999). Consequently, Gαo should not be regarded as an exclusively vomeronasal marker when examined within the brain.

### General neuronal and axonal markers

Once the accessory region had been identified by its topography, lectin-binding profile, and selective Gαi2 expression, NCAM, NSE, NeuN, PGP 9.5, and β-tubulin III provided complementary evidence concerning its neuronal composition. These markers are not specific to the vomeronasal pathway, but their convergent expression demonstrated that the identified region contained differentiated neurons, sensory projections, and organized neuropil rather than representing an isolated bundle of afferent fibers or an extension of the surrounding white matter (Thompson et al. 1983; Taniguchi et al. 1993a; Roskams et al. 1998).

Within the AOB, NCAM immunoreactivity was concentrated in the superficial glomerular-like structures previously identified using UEA, LEA, and STA. Neural cell adhesion molecules participate in sensory axon fasciculation, targeting, and glomerular organization within the olfactory system (Schwanzel-Fukuda and Pfaff 1994; Treloar et al. 2003). The correspondence between NCAM-positive profiles, lectin-reactive structures, and Gαi2-positive glomerular domains therefore supports their interpretation as accessory sensory afferents and terminal fields. NeuN provided direct evidence for the neuronal nature of the accessory region. Its immunoreactivity was detected in the nuclei of the large cells interpreted as AOB projection neurons, as well as in smaller surrounding neuronal populations. NSE showed a broader pattern involving neuronal elements and neuropil, whereas PGP 9.5 immunoreactivity was predominantly associated with the neuropil, although individual neuronal somata were also labeled. β-tubulin III strongly labeled the principal cells and their processes, revealing an organized cytoskeletal architecture despite the absence of conventional lamination.

In the MOB, the same markers revealed the organization of a much more clearly stratified neuronal network. NCAM immunoreactivity was prominent in the superficial layers and encompassed the glomerular population, although labeling intensity varied among individual glomeruli. This heterogeneity may reflect molecular differences among olfactory sensory projections and their terminal fields, as previously reported for neural cell adhesion molecules in the mammalian MOB (Treloar et al. 2003; Treloar et al. 2010). NSE broadly labeled mitral, tufted, periglomerular, and granule cells, whereas the superficial olfactory nerve layer remained largely negative. NeuN similarly identified multiple neuronal populations distributed throughout the bulb. PGP 9.5 labeled sensory afferent fibers together with mitral, tufted, and periglomerular neurons, consistent with its established value as a neuronal marker in mammalian olfactory tissues. β-tubulin III provided particularly clear visualization of MOB projection neurons. Mitral and tufted cells displayed strongly immunoreactive somata and processes, while the external plexiform layer showed intense labeling corresponding to the high density of dendritic profiles within this compartment. By contrast, the granule cell layer was weakly immunoreactive. This pattern reflects the markedly compartmentalized cytoskeletal organization of the MOB and is consistent with the differentiation of projection-neuron dendrites and local bulbar circuitry described in other mammals (Roskams et al. 1998; Tufo et al. 2022; Ruiz-Rubio et al. 2026).

### Differential OMP expression in the MOB and AOB

OMP expression revealed a pronounced difference between the accessory and main olfactory bulbs. Within the AOB, labeling was weak and restricted to a small number of fibers, with no clear immunoreactivity in the glomerular-like structures identified by UEA, LEA, NCAM, and Gαi2. This limited expression may reflect a relatively low density of vomeronasal afferents, heterogeneous OMP expression among vomeronasal sensory neurons, or a distinct molecular phenotype of the accessory projections. A comparable dissociation has been reported in both the capybara and meerkat, in which OMP strongly labels the nerve and glomerular layers of the MOB but produces only very limited labeling in the AOB, despite the advanced morphological and neurochemical maturity of their vomeronasal system (Torres et al. 2020; Torres et al. 2021). Regional heterogeneity in OMP expression has also been demonstrated in the gray short-tailed opossum (*Monodelphis domestica*), whose rostral AOB is markedly more immunoreactive than its caudal division (Shnayder et al. 1993). Because OMP expression varies among mammalian species and sensory neuronal populations, weak labeling should not be interpreted as evidence that the AOB is inactive. Rather, it indicates that the molecular phenotype -and possibly the maturation state or relative composition-of its afferent component differs substantially from that of the main olfactory pathway.

By contrast, the MOB displayed intense OMP immunoreactivity in the olfactory nerve and glomerular layers, consistent with its expression in mature olfactory sensory neurons and their central axonal projections. Not all glomeruli were labeled with the same intensity, with strongly immunoreactive structures intermingled with weakly labeled or nearly negative glomeruli. This heterogeneous distribution may reflect differences in axonal density, sensory-neuron maturation, turnover, or molecular identity among olfactory afferent populations (Farbman and Margolis 1980). The marked contrast between intense OMP expression in the MOB and its restricted distribution in the AOB provides additional evidence that the two structures receive molecularly distinct sensory projections.

### Calcium-binding proteins and neuronal specialization

Calretinin represented one of the most informative markers for differentiating the neuronal organization of the AOB. Strong immunoreactivity was detected in both its glomerular-like structures and large presumptive projection neurons. The expression of calretinin by these principal cells distinguished them from MOB mitral cells and provided an additional neurochemical criterion for delimiting the accessory region. Calretinin-positive projection neurons have been described in the AOB of other mammals, although their abundance and distribution vary substantially among species (Jia and Halpern 2004; Torres et al. 2020; Torres et al. 2021). Their prominence in *A. cinereus* may therefore represent a distinctive feature of the lutrine AOB and indicates that its reduced lamination is accompanied by a specific neuronal phenotype rather than generalized neurochemical simplification.

Within the MOB, calretinin immunoreactivity was principally associated with periglomerular cells and selected neurons located in deeper layers. This pattern agrees with previous descriptions of calretinin-positive interneuronal populations in the mammalian MOB, where they participate in the modulation of glomerular and granular circuits (Kosaka and Kosaka 2007; Schwaller 2014; Torres et al. 2021). Individual glomeruli exhibited different levels of calretinin immunoreactivity, although calretinin-positive periglomerular cells were consistently present around them. This finding further supports the existence of neurochemically heterogeneous glomerular modules within the MOB.

Neither calbindin nor parvalbumin immunoreactivity was detected in the AOB. In contrast, within the MOB, calbindin labeling was mainly associated with a small population of periglomerular interneurons, whereas parvalbumin identified an even more restricted neuronal subset. These complementary patterns are consistent with the neurochemical heterogeneity of inhibitory interneurons in the mammalian MOB and indicate that calretinin-, calbindin-, and parvalbumin-positive cells represent partially distinct components of local bulbar circuitry (Andressen et al. 1993; Kosaka and Kosaka 2007).

### Genotype-phenotype decoupling and mosaic evolution

Considered together, the available genomic and anatomical evidence supports a partial decoupling between genotype and phenotype during the evolution of the lutrine vomeronasal system. Genomic studies have identified extensive pseudogenization of olfactory receptor genes in the sea otter and giant otter, while TRPC2, an essential component of canonical vomeronasal signal transduction, has been described as a pseudogene in *Lontra canadensis* (Yu et al. 2010; Beichman et al. 2019). Nevertheless, *L. canadensis* retains an anatomically recognizable VNO, and the present study demonstrates that *A. cinereus* possesses an identifiable and neurochemically differentiated AOB. Thus, the loss or reduction of particular receptor or transduction genes does not necessarily coincide with the immediate disappearance of the peripheral organ or its central target. Although genomic and anatomical variation are broadly correlated across mammalian olfactory systems (Garrett and Steiper 2014), the different components of these systems need not undergo evolutionary reduction in a strictly synchronous manner. Instead, receptor repertoires, intracellular transduction pathways, peripheral sensory epithelia, neural projections, and central processing structures may respond differently and at different rates to ecological transitions (Hecker et al. 2019; Palmer et al. 2026). This pattern is consistent with a mosaic model of chemosensory evolution and cautions against classifying the vomeronasal system simply as either fully preserved or entirely lost. Individual components may remain anatomically recognizable while undergoing varying degrees of molecular, neurochemical, or cytoarchitectural reorganization. The combination observed in *A. cinereus* -a prominent, conventionally laminated MOB together with a small and poorly laminated but neurochemically differentiated, Gαi2-positive AOB-provides a compelling neuroanatomical example of this evolutionary mosaicism.

The strong Gαi2 immunoreactivity of the AOB is particularly relevant in this context because it indicates preservation of a molecular component commonly associated with the V1R pathway. However, neither Gαi2 expression nor the anatomical presence of an AOB alone demonstrates that the entire canonical vomeronasal transduction cascade remains operational. Conversely, pseudogenization of a single component, such as TRPC2, does not necessarily imply the immediate developmental disappearance of all peripheral and central structures associated with the pathway. Species-specific genomic and transcriptomic analyses of *A. cinereus* will therefore be necessary to establish how its preserved neuroanatomical phenotype relates to the condition of its vomeronasal receptor and transduction-gene repertoires (Yu et al. 2010; Hecker et al. 2019; Palmer et al. 2026).

### Anatomical presence versus functional preservation

These findings also illustrate the need for greater precision when describing a vomeronasal system as functional, vestigial, rudimentary, or reduced. Anatomical preservation of a VNO, vomeronasal groove, or AOB does not by itself demonstrate sensory functionality. Conversely, small size, poor lamination, or a reduced receptor repertoire does not establish complete loss of the pathway. Evidence of functionality should ideally integrate the presence of sensory neuroepithelium, receptor and transduction molecules, vomeronasal nerve projections, synaptic organization within the AOB, connectivity with higher-order targets, and physiological or behavioural responses to chemical stimulation.

In *A. cinereus*, the convergence of topographical evidence, UEA binding, Gαi2 immunoreactivity, NCAM-positive superficial structures, and differentiated neuronal populations provides substantially stronger evidence than anatomical location alone. It demonstrates that this region represents a preserved and molecularly differentiated accessory olfactory relay. The most precise interpretation is therefore that the vomeronasal pathway is anatomically retained and neurochemically differentiated, rather than that its complete sensory functionality has been experimentally demonstrated.

No single anatomical or molecular feature would have been sufficient to identify the AOB of *A. cinereus* unequivocally. Its recognition instead rests on several independent and convergent criteria: the expected dorsocaudal position within the olfactory peduncle; irregular glomerular-like regions and presumptive projection neurons revealed by Nissl staining; selective UEA binding and complementary LEA and STA profiles; strong Gαi2 immunoreactivity; and expression of NCAM, NSE, NeuN, PGP 9.5, β-tubulin III, and calretinin. This convergence establishes the accessory identity of the region despite the absence of conventional lamination.

From an ecological perspective, retention of both olfactory systems may be particularly relevant in a socially complex mustelid alternating between aquatic and terrestrial environments. Volatile cues processed by the main olfactory system may operate over the air–substrate interface, whereas less volatile components of urine, faeces, or glandular secretions deposited at latrines could remain accessible during direct nasal investigation. Recurrent use of latrine sites by Asian small-clawed otters supports the importance of persistent chemical information in their social ecology. The coexistence of a highly developed MOB and a reduced but differentiated AOB may therefore represent complementary processing of chemically diverse signals rather than simple replacement of the accessory pathway by the main olfactory system.

In conclusion, the olfactory bulbs of *Aonyx cinereus* exhibit a markedly asymmetric organization. The MOB is prominent, canonically laminated, and composed of well-differentiated projection neurons and interneuronal populations, consistent with preservation of a highly developed main olfactory pathway. In contrast, the AOB is small, macroscopically indistinguishable, and lacks the layered architecture characteristic of classical rodent models. Nevertheless, its topographical position, neuronal composition, selective UEA binding, strong Gαi2 immunoreactivity, and distinctive calretinin expression demonstrate that it constitutes a genuine and neurochemically differentiated accessory olfactory relay.

Rather than representing either complete preservation or complete regression, the vomeronasal system of lutrines appears to exhibit a mosaic organization in which genomic, peripheral, and central components have followed partially independent evolutionary trajectories. Together with the recent identification of the VNO in *Lontra canadensis*, the present findings provide evidence for persistence of peripheral and central vomeronasal components in semiaquatic mustelids. *Aonyx cinereus* therefore represents an informative comparative model for investigating how phylogeny, social communication, and secondary adaptation to aquatic environments have shaped the evolution of mammalian olfactory systems.

## AUTHORS CONTRIBUTION

Silvia Alejandra FRAGA-ABELLEIRA: Conceptualization; Methodology; Investigation; Writing - original draft; Writing - review & editing. Mateo V. TORRES: Conceptualization; Investigation; Methodology. Rocío CANALES: Investigation; Methodology. Sara RUIZ-RUBIO: Conceptualization; Investigation; Methodology. Pablo SANCHEZ-QUINTEIRO: Conceptualization; Investigation; Methodology; Funding acquisition; Writing - original draft; Writing - review & editing; Supervision. Irene ORTIZ-LEAL: Conceptualization; Investigation; Methodology; Writing - review & editing; Methodology; Supervision.

## COMPLIANCE OF ETHICAL STANDARDS

### Ethical approval

The specimens examined in this study were obtained from Asian small-clawed otters (*Aonyx cinereus*) housed at a zoological institution that died of natural causes unrelated to the present research. No animals were euthanized or subjected to experimental procedures for the purposes of this study. Consequently, specific ethical approval for animal experimentation was not required under Directive 2010/63/EU and applicable national legislation. The collection and scientific use of the post-mortem material were authorized by the zoological institution.

### Informed consent

No human subject was used in this study.

### Data availability

Not Applicable.

